# Microbiome composition shapes rapid genomic adaptation of *Drosophila melanogaster*

**DOI:** 10.1101/632257

**Authors:** Seth M. Rudman, Sharon Greenblum, Rachel C. Hughes, Subhash Rajpurohit, Ozan Kiratli, Dallin B. Lowder, Skyler G. Lemmon, Dmitri A. Petrov, John M. Chaston, Paul Schmidt

## Abstract

Population genomic data has revealed patterns of genetic variation associated with adaptation in many taxa. Yet understanding the adaptive process that drives such patterns is challenging - it requires disentangling the ecological agents of selection, determining the relevant timescales over which evolution occurs, and elucidating the genetic architecture of adaptation. Doing so for the adaptation of hosts to their microbiome is of particular interest with growing recognition of the importance and complexity of host-microbe interactions. Here, we track the pace and genomic architecture of adaptation to an experimental microbiome manipulation in replicate populations of *Drosophila melanogaster* in field mesocosms. Manipulation of the microbiome altered population dynamics and increased divergence between treatments in allele frequencies genome-wide, with regions showing strong divergence found on all chromosomes. Moreover, at divergent loci previously associated with adaptation across natural populations, we found that the more common allele in fly populations experimentally enriched for a certain microbial group was also more common in natural populations with high relative abundance of that microbial group. These results suggest that microbiomes may be an agent of selection that shapes the pattern and process of adaptation and, more broadly, that variation in a single ecological factor within a complex environment can drive rapid, polygenic adaptation over short timescales.

**Significance statement:** Natural selection can drive evolution over short timescales. However, there is little understanding of which ecological factors are capable of driving rapid evolution and how this rapid evolution alters allele frequencies across the genome. Here we combine a field experiment with population genomic data from natural populations across a latitudinal gradient to assess whether and how microbiome composition drives rapid genomic evolution of host populations. We find that differences in microbiome composition cause divergence in allele frequencies genome-wide, including in genes previously associated with local adaptation. Moreover, we observed concordance between experimental and natural populations in terms of the direction of allele frequency change, suggesting that microbiome composition may be an agent of selection that drives adaptation in the wild.

## Introduction

A growing number of studies have identified genes that contribute to adaptation (1–4), but the ecological mechanisms that drive evolution are rarely identified (5). Ecological factors often co-vary in nature, so disentangling the effects of putative agents of selection on changes in allele frequencies requires experimental manipulation. Patterns of intraspecific genomic variation in nature can be shaped by differences in founder populations, connectance between population, and demography, complicating inferences of selection (6). Replicated selection experiments provide a way to test whether particular ecological mechanisms act as agents of selection and assess the genomic architecture of adaptation, both key challenges to understanding adaptation (2, 6–8). Yet, using selection experiments to identify mechanisms capable of driving rapid evolution in nature also presents methodological challenges; it is difficult to create both ecologically realistic (e.g. complex selective environment, population sizes allowed to varying across treatments) and evolutionarily realistic (e.g. sufficient standing genetic variation, multiple generations, selection agents similar to those in nature) conditions that allow experimental results to translate to populations in nature (5). Combining field selection experiments with population genomic data from both experimental and natural populations presents a powerful approach to determine whether and how particular agents of selection drive rapid evolution in the genome.

Many prominent theories in evolution suggest that species interactions are the primary mechanism that drives evolution and diversification (9–14). Yet, determining which species interactions actually drive rapid evolution when selective landscapes are complex is crucial to understanding both the mechanisms and outcomes of adaptation (15–17). Outdoor experiments that manipulated specific species interactions have provided convincing evidence that competition and predation can act as agents of selection capable of driving rapid phenotypic evolution (18–21). Host-microbe interactions can be strong and there is evidence they can drive macroevolutionary patterns (22–26), but associated microorganisms have not been experimentally investigated as an agent capable of driving rapid host evolution (27, 28) except where symbiont evolution is tied to the host through vertical transmission (29, 30). Bacteria play a crucial role in the physiology, ecology, and evolution of animals even if they are not transmitted or acquired across generations (22, 31–34) and the composition of affiliated microbial communities can impact host performance and relative fitness (35). Moreover, patterns of intraspecific variation in microbiome composition that could have considerable effects on host physiology and performance have been described in a growing number of taxa (36–39). The amount of intraspecific variation in microbiome composition and its effects on host phenotypes have led to considerable speculation, but little data, on the important role the microbiome may play in host evolution (27, 28, 34, 40).

*Drosophila melanogaster* presents an excellent system in which to investigate whether microbiome composition acts as an agent that drives rapid host genomic adaptation. *D. melanogaster* populations vary in their microbiome composition in eastern North America, driven by latitudinal variation in the relative proportion of acetic acid bacteria (AAB) and lactic acid bacteria (LAB) (41). Inoculation experiments in the lab have demonstrated that LABs and AABs directly influence the functional traits of *D. melanogaster* including development rate, lipid storage, and starvation tolerance (42, 43). *D. melanogaster* populations in eastern North America have long been a model for testing hypotheses of local adaptation, as there are strong patterns of both phenotypic and genomic evolution across latitudes that covary with temperature and photoperiod (44–48). Extensive genomic sequencing of natural populations has revealed thousands of independent SNPs that vary clinally and hence are likely involved in adaptation (46, 48). Finally, large *D. melanogaster* populations can be manipulated in replicated field mesocosms providing the opportunity to connect the wealth of genomic information about this species with an understanding of evolution in natural contexts.

To test whether microbiome composition can drive rapid evolution we introduced outbred populations of *D. melanogaster* into 14 individual 2m × 2m × 2m outdoor experimental enclosures. We then applied three treatments to these populations as they evolved over a 45 day period: 1) addition of the AAB species *Acetobacter tropicalis* to the food resource (*At* treatment) 2) addition of the LAB species *Lactobacillus brevis* to the food resource (*Lb* treatment) 3) no microbial inoculation (*No-Ad* treatment). We used 16s rRNA sequencing and microbial culture to ascertain the efficacy of the treatments and tracked host population size in each replicate to determine whether treatments altered host population dynamics. We tested for rapid evolution in response to microbiome treatments by coupling whole genome data for each replicate with previously identified lists of putatively adaptive loci and examining whether microbiome treatments led to enhanced genomic divergence relative to control populations. In addition, we compared the direction of allele frequency change to determine whether differences between experimental treatments were similar to those observed in natural populations as a way of assessing the importance of microbial variation in driving adaptation across natural populations.

## Results and Discussion

### Efficacy of shifting the microbiome in an outdoor experiment

Microbial addition treatments shifted the overall microbiome composition of *D. melanogaster* populations (Bray Curtis F_1,29_= 15.8, p<0.001, Fig. 1A, Unifrac metrics in Fig. S1) and the relative abundance of individual operational taxonomic units (OTUs) and the abundance of colony forming units (CFUs) (Figures S2, S3, and S4). While the different treatments displayed substantial variation in the relative abundance of AAB and LAB, both microbial groups were present in the microbiome of all experimental populations (Fig. S3). Sequencing the V4 region of the 16S rRNA gene demonstrated that microbiomes of *D. melanogaster* in *At*- and *Lb*-treated cages were enriched for OTUs with perfect identity to the 16s rRNA gene of *At* and *Lb*, respectively. In addition, whole genome sequencing of randomly selected microbial colonies isolated from one *At*-treatment cage revealed AAB with >99.9% whole-genome similarity to the added *At* strain (Fig. S4). Overall, the differences in microbiome composition between the *At* and *Lb* treatments are modest compared to population-level differences in microbiome composition found across latitudes, where high-latitude locations have microbiomes dominated by LAB and microbiomes in low-latitude populations are dominated by AAB (41).

**Figure 1:**
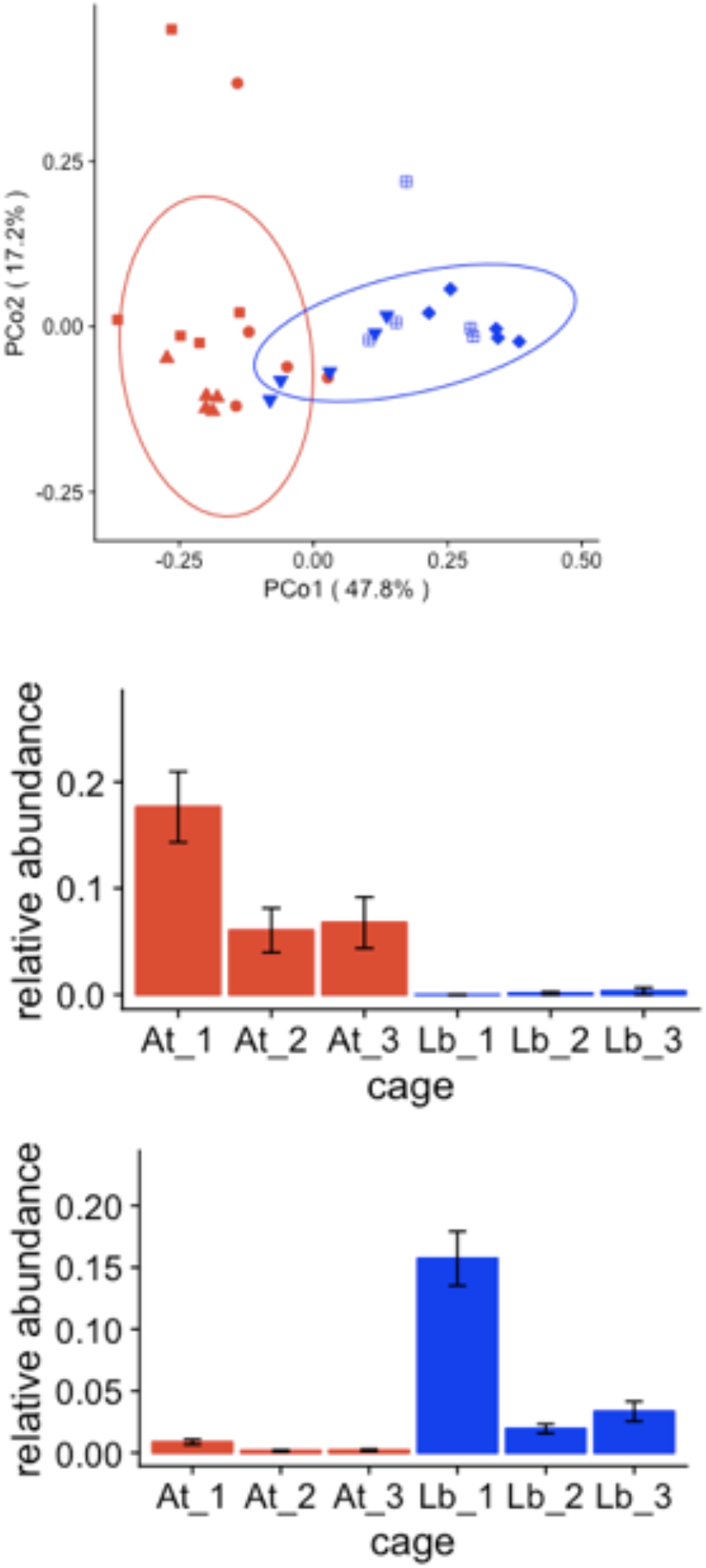
The effect of microbial additions on the gut microbiomes of *D. melanogaster* in the *At* and *Lb* treatments. Panel A shows the effect of *At* and *Lb* treatments at the fourth week of the experiment on microbiome composition of pools of adults males collected from cages. Panels B and C show the relative abundance of AAB and LAB (respectively) in the microbiomes of *D. melanogaster* from each microbial addition treatment (plotted as means +/− SEM).

The influences of distinct AAB and LAB on various *D. melanogaster* phenotypes are well characterized (42, 49–53). To test whether previously reported phenotypic effects are also detectable in outbred *D. melanogaster* populations we compared the larval development of individuals from the *No-Ad* experimental cages when monoassociated with *At* and *Lb*. Consistent with previous work, bacterial treatment significantly influenced larval development time: *At* led to ~10% higher development rate than *Lb* (Z=-15.9, P<0.001). The effects of microbiome composition on host ecology presents a general mechanism by which microbiomes may shape rapid evolution of host populations.

### Influences of microbiota treatments on host ecology

To determine whether microbiome communities alter the ecological characteristics of host populations in outdoor mesocosms, and hence could plausibly shape host evolution, we measured two key ecological characteristics in field mesocosms: fly body mass and population size. Individuals collected directly from *At* treatment populations had 28% higher mass than those from *Lb* treated populations (F_2,19_=13.81, p=0.0002) (Fig. 2A). We also observed increased sexual dimorphism in *At* treatments in body size relative to the *Lb-* and *No-Ad* treatments (F_2,19_=5.73, p=0.0113). In contrast, *Lb* replicates had significantly higher population sizes than *At* replicates (chisq=14.86, df=1, p=0.0001, Fig. 2B) suggesting that microbiome treatments influence the tradeoff between somatic and reproductive investment. The difference in population size demonstrates that shifts in the relative abundance of the *D. melanogaster* microbiota can significantly alter host population dynamics. Differences in population size associated with microbiome composition provides clear evidence to support previous assertions that natural population-level variation in the microbiota that has been observed across the animal kingdom (39, 41, 54, 55) may influence the population ecology of hosts bearing diverse communities of partners (28, 34, 56). Such patterns are established for hosts bearing obligate partners (57–59) or infected with microbial symbionts (60), but our data demonstrate that changes in the relative abundance of microbial taxa can shape host populations. These differences in body size and population dynamics, due to a presumed combination of ecological and evolutionary forces, demonstrate that modest shifts in microbiomes can alter host populations in natural settings which bolsters the hypothesis that microbiomes could drive rapid evolution.

**Figure 2:**
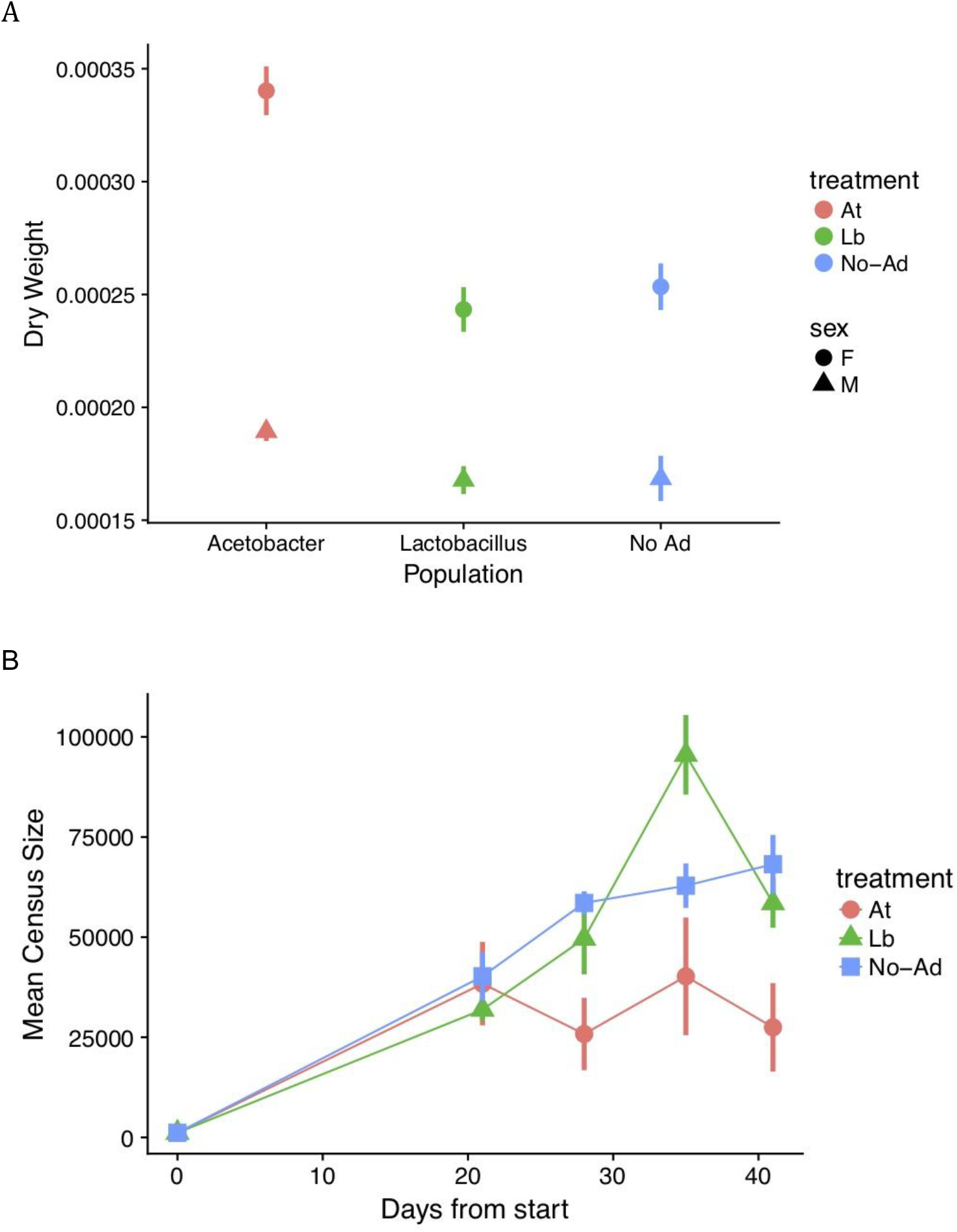
Population size and body mass of *D. melanogaster* populations from each microbial addition treatment. Panel A shows shows the mean from each treatment of the dry weight of *D. melanogaster* individuals of each sex from each replicate cage. Panel B shows host population size over the course of the experiment. Both panels values plotted are means +/− SEM.

### Microbiome composition shapes host genomic evolution

We assessed whether differences in microbiome composition across *At* and *Lb* treatments shaped *D. melanogaster* evolution over the course of five host generations. Using a whole genome pool-seq approach (61), we generated data on allele frequencies at 1,988,853 biallelic segregating sites after filtering (see Methods) for the founder population and from each experimental replicate after 45 days of microbiome treatment. Given that our experiment was founded with a genetically diverse population with little linkage disequilibrium (62) and any divergent selection between treatments was limited to 5 overlapping generations, we did not expect substantial genome-wide divergence (63, 64). To assess any genome-wide divergence, we calculated the mean F_ST_ statistic between the founder population and the three treatment populations, for subsets of 1,000 sites sampled randomly from across the genome (Fig. S6). We also conducted a principal component analysis of allele frequencies from all sampled populations to visualize divergence genome-wide (Fig. S7). In both analyses we observe a trend that microbial treatment (both *At* and *Lb*) prompts greater genome-wide divergence from the founder population than *No-Ad* over the relatively short duration of the experiment. We also assessed divergence between treatments in smaller overlapping windows of the genome and found significantly enhanced divergence between pairs of *At* and *Lb* treated cages compared to pairs of *No-Ad* cages (p<2.2e-16, Welch’s two-sample t-test) (Fig. 3). Signatures of this enhanced divergence across microbial treatments were observed across the genome and on all chromosomes. This pattern of enhanced divergence due to microbial differences demonstrates that modest variation in microbiome composition can drive genomic divergence of host populations over short and ecologically relevant timescales.

**Figure 3:**
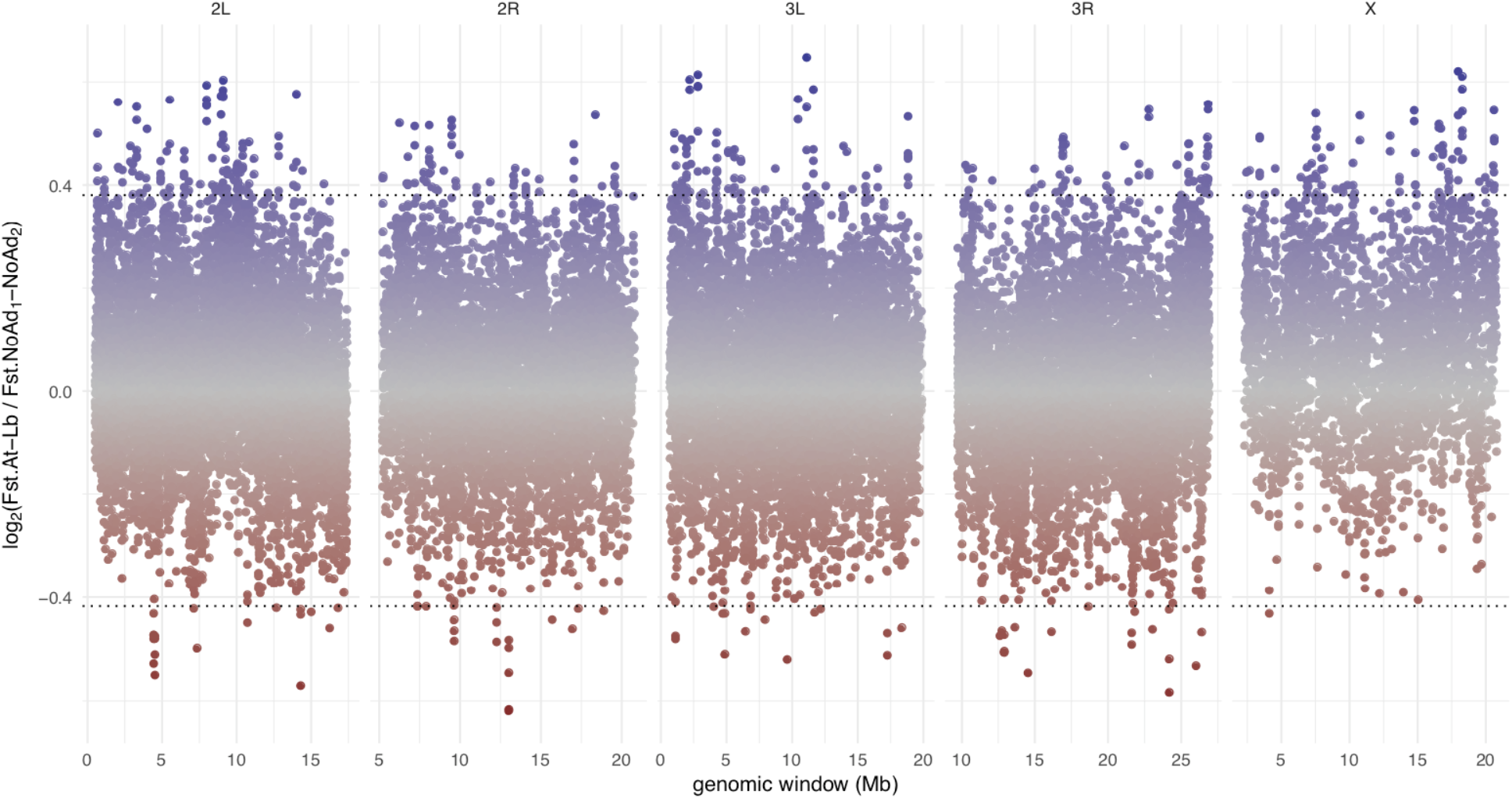
Genomic landscape of divergence between *At* and *Lb* treated cages compared to divergence observed amongst *No-Ad* cages. Fst was calculated between pairs of samples in windows of 250 SNPs, tiled across the genome with 50-SNP shifts. Shown below is the log2 of the ratio between average Fst between pairs of *At* and *Lb* samples compared to average Fst between pairs of *No-Ad* samples for each window (positive values show more divergence between *At* and *Lb*, negative more divergence among *No-Ad*). Panels are chromosomes and the black dotted lines show the values for 2.5 standard deviations above and below the mean. The enrichment for values >2.5 standard deviations above the mean relative to those below the mean demonstrate enhance divergence between *At* and *Lb* treatments. Centromeric and telomeric regions were excluded from this analysis according to the coordinates provided in (90).

In addition to whole genome and window-based analyses we also assessed patterns of divergent selection between *At* and *Lb* populations at individual sites. Linkage disequilibrium decays over ~200bp in most regions of the the *D. melanogaster* genome (62) and our founding populations contained substantial standing genetic variation, giving us considerable genomic resolution with which to detect selection. To assess divergent selection between treatments at each segregating site we fit a generalized linear model to allele frequencies as a function of microbiome treatment, accounting for replicate cage as an independent factor. We found 297 sites diverged significantly between *At* and *Lb* treatments with FDR<.05 and minimum effect size of 2% (Table S1). These sites were located on all chromosomes and were found in or near 281 genes, indicating little linkage between significant sites. The *D. melanogaster* genome contains several inversions that vary in frequency across populations in a way that is suggestive of adaptation (65), but we observed no enrichment for divergence of inversion frequencies associated with microbial treatment (based on marker sites, Table S2), meaning overall patterns of divergence were not driven by shifts in inversion frequencies. The pattern of divergence we observed across resolutions, both at individual sites and in an analysis based on small windows, demonstrates that the genomic response to microbiome treatments has a complex genetic architecture, with signatures of selection at many independent regions of the genome. These results fit with a polygenic model of adaptation, in which many genes contribute to adaptation (66), and suggest that the genomic basis of adaptation over very short timescales can be polygenic.

### Links between microbiome manipulation and changes in allele frequency in nature

Combining our experiment with population genomic data from nature allows us to test whether differences in microbiome composition alone are capable of driving divergence in allele frequencies at SNPs that vary across natural populations. Along the east coast of North America, high-latitude populations of *D. melanogaster* have LAB-enriched microbiomes and populations from lower latitudes have AAB-enriched microbiomes (41). Comparative genomics work has identified sites that are likely adaptive along this cline (67), 15,399 of which varied in our experimental populations. We tested whether the allele that was more common in populations experimentally enriched for a microbial group was also more common in the natural clinal population that has a high relative abundance of the same microbial group. We labeled sites as ‘directionally concordant’ if the allele that was at higher frequencies in high-latitude populations compared to low-latitude populations was also the allele that was at higher frequencies in *Lb* populations compared to *At* populations. When we considered all ~2 million variant sites, the percent of directionally concordant sites was 50.3%, indistinguishable from a null expectation. However, concordance rose significantly in subsets of sites with both strong divergence between microbial treatments and strong clinal variation (Fig. 4). For example, 70.7% were concordant among the 945 SNPs with *At-Lb* divergence pval<.05, effect size>2%, and clinal p-value<10^−5^, while 80.0% were concordant among the 35 SNPs with *At-Lb* divergence pval<.01, effect size>2%, clinal p-value<10^−8^. 1,000 rounds of randomly sampling sites matched to observed data for chromosome and allele frequency demonstrated that these concordance values are both significantly higher than expected by chance (p<0.001 in both cases). In the latter case, the majority of the 35 SNPs are on chromosome arm 3R, yet are located in or near 32 different genes, several of which are known to play a role in local adaptation (67–69) (Table S3). Though these high levels of concordance at top divergence sites may suggest long-range linkage disequilibrium, we did not find significantly elevated concordance in any of 7 large chromosomal inversions (Table S2). The surprising concordance of the identity of AAB-associated and LAB-associated alleles in experimentally-treated populations and natural clinal populations suggests microbiome composition may be a significant component of the fitness landscape, and hence adaptation, in natural populations.

**Fig. 4:**
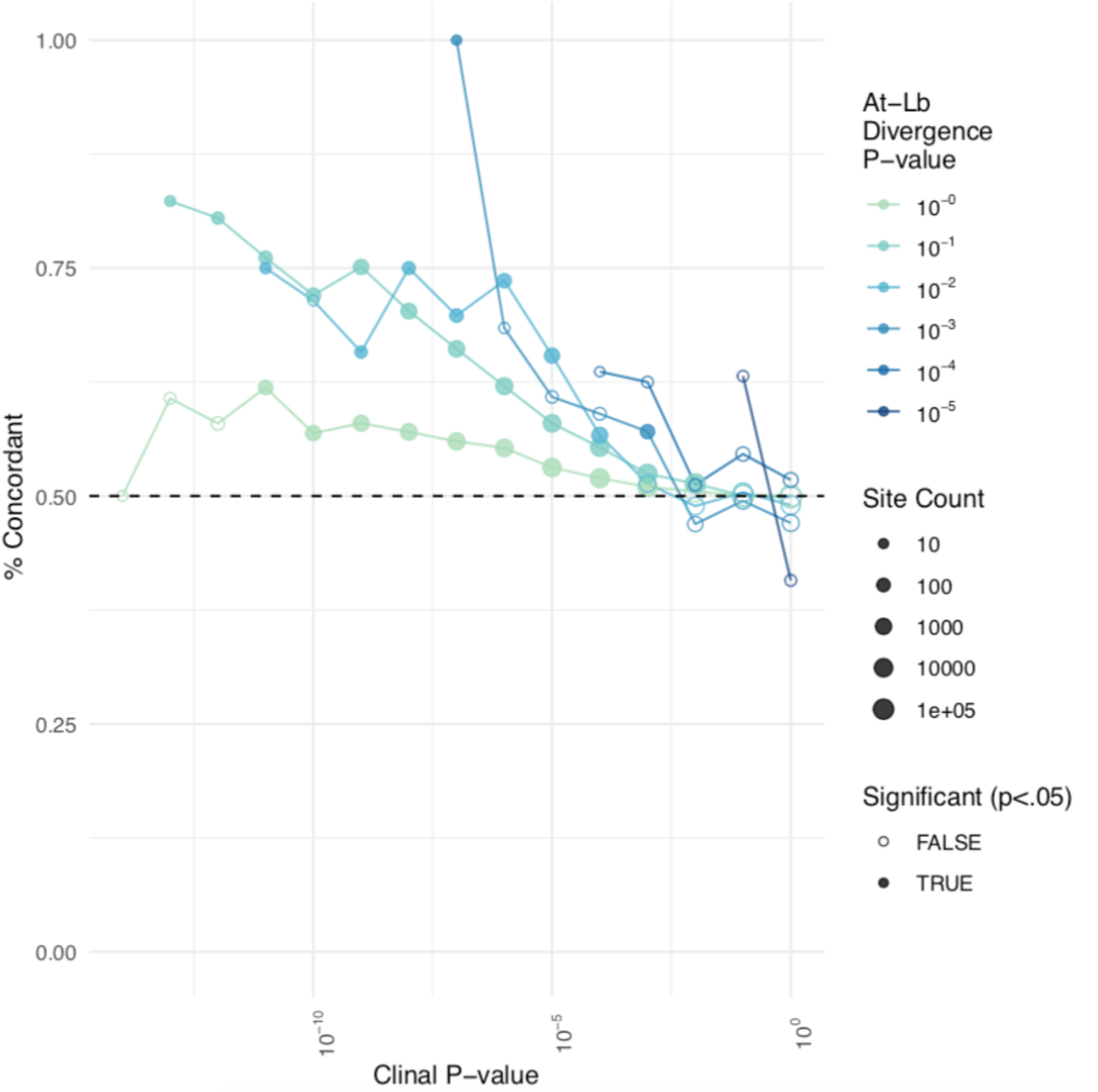
Concordance of allelic divergence in natural and experimental populations. Concordance is calculated as the % of sites in which the allele found at higher frequencies in natural high-latitude populations compared to low-latitude populations was also found at higher frequencies in experimental *Lb* populations compared to *At* populations. Each point refers to a distinct subset of sites, binned according to clinality (x-axis) and *At*-*Lb* divergence (color); the number of sites examined is indicated by the size of the point. A dashed black line is drawn at the null expectation of 50% concordance. Solid-colored points represent site subsets in which concordance is significantly elevated compared to the shuffled null distribution.

## Conclusion

Moving from documenting cases of rapid evolution to studying the driving mechanisms is crucial to understanding adaptation in natural populations (16). Microbiomes can influence nearly all aspects of host biology (27, 40, 70) and it has long been assumed that microbiomes are also an important factor at the population-level (28, 71). Our manipulative experiment demonstrates that changes in the relative abundance of individual members of the *D. melanogaster* microbiome are sufficient to enhance genomic divergence of host populations over only 5 generations. The magnitude of divergence was heterogeneous across the genome, but we uncovered regions of strong divergence on all chromosomes. Genomic patterns also illustrate that variation in microbiome composition is a sufficiently strong agent of selection to drive evolution at loci that exhibit putatively adaptive patterns across populations in nature. We detected concordance in the directionality of allelic change at these sites between our experiment and natural populations, which provides evidence that variation in microbiome composition is a substantial component of the fitness landscape. Overall, our results demonstrate that shifts in microbiome composition can be important drivers of ecological and evolutionary processes at the population level and that a single ecological factor within a complex environment can drive polygenic adaptation over short timescales.

## Author Contributions

PS and JMC conceived of the experiment. SMR, RH, JMC and PS conducted the experiment with help from SR and OK. SMR collected population size and body size data. RH, SGL, DLL, and JMC sequenced and analyzed the microbiome and measured phenotypic evolution. SG, SMR, and DAP analyzed the genomic data. SMR wrote the initial manuscript with assistance from SG and JMC. DAP, JMC, and PS discussed the results and implications and commented on the manuscript at all stages.

## Materials and Methods

### Experimental setup

We constructed the founding *Drosophila melanogaster* population for this experiment by crossing 150 wild-collected isofemale lines from Pennsylvania. 10 males and 10 females were taken from each line and combined into a single breeding cage. After 3 generations of mating and density controlled rearing in favorable lab conditions we introduced 500 females and 500 males of a single age cohort into each experimental cage on June 15th, 2017. Subsamples of the founding population were collected on June 15th for initial genomic sequencing. Flies were in enclosures from June 15th to August 3rd 2017, which, based on larval development rates in outdoor cages allowed for ~five overlapping generations. Outdoor cages are 2m × 2m × 2m enclosures constructed of fine mesh built around metal frames (BioQuip PO 1406C) (72, 73). Inside of these enclosures we planted 1 peach tree and vegetative ground cover to provide shading and physically mimic the natural environment. Peaches were removed before ripening to prevent flies from feeding on them. Photographs of eight quadrats within each cage were taken and flies were counted to estimate population size at five time points during the experiment. We tested for effects of microbiome treatments on host population size using an LME with microbial treatment as a fixed effect and sample date as a random effect. Each cage was used as a statistical replicate and our analysis was conducted on all census data after the initial population expansion (> day 21 of the experiment).

### Microbial treatments

The experiment consisted of three treatments: diet supplemented with *Lactobacillus brevis* DmCS_003 (*Lb*), diet supplemented with *Acetobacter tropicalis* DmCS_006 (*At*), and no bacterial addition (*No-Ad*). To prepare the bacterial inoculum, a 24-72 h culture of each species was centrifuged for 10 min at 15,000× g and resuspended in phosphate buffered saline (PBS) at OD600=0.1. Separately, 300 ml of modified Bloomington diet was prepared in a 1.5lb aluminum loaf pan under standard lab conditions (non-sterile). Within 24 h of diet preparation, 2.2 ml normalized bacteria were spread on the surface of the food inside of the loaf pan. The inoculated diets were covered for a 12-36 h incubation at 25°C and transported to the outdoor experiment site 3 times each week. Diets were uncovered immediately after introduction to outdoor fly enclosures. Diets were left undisturbed for 2-3 days, and then covered with mesh caps to permit larval development but exclude egg laying adults. Caps were removed when flies had eclosed to permit release of the next adult generation into the enclosure. The protocol for the No-Ad replicates mimicked the above but did not include any inoculation of the food. The diets provided the only source of food available that was capable of supporting *D. melanogaster* development.

### Quantification of microbial communities from experimental treatments

For culture-dependent analysis, five pools of five male flies were collected from each treated outdoor cage and homogenized in a microcentrifuge tube containing 125μl mMRS medium. Homogenates were dilution plated onto mMRS and grown at 30°C under ambient and restricted oxygen conditions. Tan- or copper-colored colonies were classified as AABs, and white or yellow colonies were classified as LABs. 1ml of the same homogenate was pelleted for DNA extraction via the QuickDNA Fecal/Soil Microbe kit (Zymo Research, D6011) and analyzed by culture-independent analysis as described below. Pairwise comparisons between absolute colony-forming unit (CFU) abundances were determined by a Dunn test.

We used 16S rRNA marker genes of pooled whole-body flies to survey the microbial community associated with the pooled fly homogenates. From each DNA extraction, the V4 region of the 16S rRNA gene was amplified as described previously, except using a HiSeq 2500 at the BYU DNA sequencing center (74). Sequence variants were clustered and assigned to the sequencing data using QIIME 2 (75, 76). After taxonomic assignment, sequences identified as *Wolbachia* were removed, and the OTU tables were rarefied to balance sequence depth with sample retention. The single OTUs with perfect matches to the *At* and *Lb* genomes were identified using BLASTn (77). Tests for significant differences in microbial beta-diversity (Bray-Curtis, weighted Unifrac, unweighted Unifrac) were performed in R using PERMANOVA (78). Differences in taxonomic abundance were assessed using ANCOM, which uses relative abundances to assess differences in community composition (79). Figures were created using ggplot2 (80).

### Measuring body size and development rate

At the conclusion of the experiment we sampled adult individuals from all cages. To determine adult mass content of cage-caught individuals, we took pools of five individuals of each sex, dried them at 55c for 24 hours, weighed them, and divided the total weight by five to obtain average individual mass. Body size data (dry weight) were analyzed using a ANOVA with microbial treatment and sex as fixed effects with cage used as the unit of replication.

We collected eggs from each *No-Ad* cage to determine the effect of monoassociation with *At* and *Lb* on development rate. To rear in monoassociation, fly eggs were collected within 24 h of deposition, bleached twice for 150s each, rinsed thrice in sterile H_2_O, transferred to sterile diet at a target density of 30-60 eggs per vial, and inoculated with a phosphate-buffered saline (PBS)-washed overnight culture of either bacterial species, normalized to OD_600_= 0.1 (81). The period of larval development was determined by counting the number of empty pupae in each vial three times each day (at 1, 6, and 11 hours into the daily light cycle) until all flies had eclosed or until no flies eclosed in three consecutive time periods, whichever came first. Bacteria-dependent differences in *D. melanogaster* development were analyzed using Cox mixed survival models in R. Development rate was calculated as the inverse time to eclosion. Significant differences between treatments were determined by a Cox proportional hazards model, analyzed separately for each bacterial inoculation, and are reported as different letters over the symbols. Summary statistics were also calculated by ANOVA.

### Genomic sequencing

We sequenced pools composed of 120 males and 80 females collected from each cage at the end of the study. We extracted the DNA and prepared libraries using ~500bp fragments for whole genome sequencing using (KAPA Hyper Prep kit). Libraries were multiplexed with dual-indexing and sequenced on multiple lanes of an Illumina NovaSeq (6 samples on each lane) system with 150bp paired end reads. Reads were checked for quality using FastQC. Adapters were trimmed with Skewer (82) and reads with a quality score <20 were removed, and overlapping read pairs were merged with PEAR (83). We aligned reads to a reference genome composed of the *D. melanogaster* reference sequence (**v5** (84)), the *Lactobacilis brevis*, and the *Acetobacter tropicalis* genomes using BWA (85), then removed duplicate reads with Picardtools and realigned remaining reads around indels with GATK’s IndelRealigner (86). Index switching, where reads are attributed to the wrong sample, can happen on Illumina HiSeq platforms (87). We detected a small amount of human contamination, likely due to index switching, and removed all reads that mapped to the human genome using bbmap, with parameters suggested at https://jgi.doe.gov/data-and-tools/bbtools/bb-tools-user-guide/bbmap-guide/ for removing contaminant reads while minimizing false positives (minratio=0.9 maxindel=3 bwr=0.16 bw=12 fast minhits=2 qtrim=r trimq=10 untrim idtag printunmappedcount kfilter=25 maxsites=1 k=14) and a version of human reference genome hg19 masked for repeat short kmers, low entropy windows, and regions highly conserved across species. This reference genome was created by Brian Bushnell specifically for human contaminant removal, and is freely available at https://drive.google.com/file/d/0B3l...it?usp=sharing, full description at http://seqanswers.com/forums/showthread.php?t=42552. After mapping and QC we retained an average of 83M mapped reads per sample at an average coverage (mosdepth (88)) of 109× of the *Drosophila melanogaster* autosomes (range 92-133×) and average coverage 92× on the X chromosome. We then used PoPoolation2 (Kofler et al. 2011) to obtain allele counts at segregating sites, discarding bases with quality <20. To be included for downstream analysis we required SNPs to be bi-allelic with one of the two alleles matching the reference allele, and we excluded SNPs overlapping any called indels, SNPs with less than 10 mapped reads containing the minor allele (an allele frequency of ~0.5% across all samples), and SNPs with min and max read depths less than 50 or greater than 250 respectively. Since the timescale of our experiment was too short to expect any true signal from new mutations arising during the 5 generations of evolution, we additionally filtered out any SNPs with allele frequencies <1% in either sample from the founder population. Finally, we excluded SNPs within repeat regions as defined by UCSC RepeatMasker (89), and any SNPs that showed distinct allele frequency ranges in the two rounds of sequencing. This yielded at dataset of ~2 million SNPs.

### PCA and Fst Analyses

Allele frequencies at each segregating site for each sample were used to conduct a principal component analysis using the R function *prcmp* with *scale=TRUE*, and the first two PCs were plotted to examine genome-wide divergence across samples visually. To obtain a more quantitative account of the divergence of populations under each treatment from the founder population, a bootstrap-Fst analysis was conducted with 1,000 rounds. In each round, 1,000 sites were randomly selected from across the genome, and Fst was calculated at each site between the average allele frequency in the two founder samples and allele frequencies averaged within treatment groups (3 of the 8 *No-Ad* samples were randomly averaged for each round to match the number of *At* and *Lb* samples). Next, a window-based analysis was used to examine divergence between treatments. Average Fst was calculated between allele frequencies from each pair of treatment samples in windows of 250 consecutive SNPs, with 50-SNP step-size between windows. For each window, the average Fst between the 3 *At* and 3 *Lb* samples was recorded, as well as the average Fst between 3 randomly selected *No-Ad* samples and 3 other randomly selected *No-Ad* samples. Windows overlapping centromeric or telomeric regions as defined by Comeran et al. (90) were excluded from this analysis, as the exceedingly low recombination rates in these regions could make them more prone to linked fluctuations across large numbers of sites.

### SNP Outlier analysis

To find SNPs that changed in frequency due to microbial treatment we used the R function *glm* to fit a generalized linear model to the allele frequencies at each SNP to test for significant associations between allele frequency and treatment. GLMs were fit using a quasibinomial error structure, as this reduces the rates of false positives relative to other significance testing protocols in genomic data (91). We identified outlier sites with significant divergence between *At* and *Lb* samples at an FDR <.05 (92), and a mean difference in allele frequency (effect size) of 2%, as this was approximately the average difference in allele frequency between treatments for all SNPs.

### Test for directional concordance with clinality

SNPs that vary across the North American latitudinal cline may reflect local adaptation (67–69, 93), and represent potential sources of adaptation to microbiome composition, which is one of many factors known to vary along this cline. Although we do not expect extensive overlap between SNPs that vary predictably along the cline and SNPs that vary predictably between treatments in our experiment (due to different segregating sites, different non-microbiome-related selective pressures, and different timescales of adaptation), we did predict that the subset of SNPs that are strongly predictable in both cases should be ‘oriented’ in the same direction ie., an allele strongly associated with natural clinal populations harboring more AAB should also be the allele associated with experimental populations experimentally enriched for AAB (here, the *At* treatment). As such, we used an existing genomic dataset on clinal variation (67, 68) to see if the SNPs that showed both 1) divergence between microbial treatments in our experiment, and 2) divergence between natural clinal populations, were more likely to be ‘directionally concordant’ than other SNPs. We first collected p-values and coefficients for each SNP in our dataset from our generalized linear model of allele frequency divergence between treatments (p_At-Lb_ and coef_At-Lb_), and p-values and coefficients from a previously conducted generalized linear model of allele frequency divergence across the cline (p_cline_ and coef_cline_). The models were oriented such that a positive coef_At-Lb_ indicated that the frequency of the alternate allele was higher in *Lt* samples than *At* samples, while a positive coef_cline_ indicated that the frequency of the alternate allele was higher in high-latitude (LAB-enriched) populations than low-latitude (AAB-enriched) populations. We assigned each SNP to two bins: an *At-Lb* divergence bin equal to the integer nearest −log10(p_At-Lb_), and a clinality bin equal to the integer nearest −log10(p_cline_). We then examined the intersection of each *At-Lb* bin and each clinality bin, and recorded the percent of SNPs where the sign of coef_At-Lb_ matched the sign of coef_cline_, which we termed ‘directional concordance’. Finally, we shuffled the bin labels across SNPs 500 times (maintaining the same bin pairs), and remeasured directional concordance values to obtain a p-value for each true concordance value.

### Tests for enrichment at inversions

We identified breakpoints (94) and segregating marker sites (95) associated with 7 large chromosomal inversions. To test for enrichment of divergence between *At* and *Lb* samples at marker sites for each inversion, we first assigned every segregating site a divergence score equal to −log10 of the p-value from the GLM analysis of per-site divergence. We then recorded the percent of times (of 1,000 replicates) that an equally-sized random set of sites had a mean divergence score higher than the markers of a particular inversion. Similarly, to test for enrichment of *At-Lb* divergence at sites within each inversion, we recorded the percent of times (of 1,000 replicates) that a randomly-selected set of 1,000 sites from outside an inversion had a mean divergence score higher than a randomly-selected set of 1,000 sites from inside an inversion. Finally, to test for enrichment of clinal concordance within each inversion, we recorded the percent of times (of 1,000 replicates) that a randomly-selected set of 1,000 sites from outside an inversion had a concordance rate higher than a randomly-selected set of 1,000 sites from inside an inversion.

## Supplemental tables and figures

**Table S1.** List of sites that show significant divergence between *At* and *Lb* cages in the experiment (effect size >2% and FDR <.05). Table shows: the chromosome arm, position, whether the SNP is within a centromeric or telomeric region, reference allele, alternate allele, gene name, p-value for divergence between *At* and *Lb* treatments, average allele frequency in founder, *No-Ad*, *At*, and *Lb* populations, p-value for divergence across clinal populations, all annotations of SNP effect according to snpEff, and GO terms.

**\*Table is large so it is attached separately.**

**Table S2.** Enrichment of At-Lb divergence among sites strongly linked to large cosmopolitan inversions (inversion markers), and enrichment of At-Lb divergence and clinal concordance among sites within inversions. In each case, the number of sites tested, and their average At-Lb divergence score is indicated, as well as a non-parametric p-value indicating the percent of times (of 1,000 trials) that a random subset of sites of equivalent size had an equal or higher average divergence score (divergence.p). For sites within inversions, the concordance rate between the identity of the allele associated with experimentally-treated and natural clinal populations enriched for the same microbial taxa is also indicated, as well as a similar non-parametric p-value.

| inversion | chrom | start | stop | inversion markers |  |  | inversion sites |  |  |  |  |
| --- | --- | --- | --- | --- | --- | --- | --- | --- | --- | --- | --- |
|  |  |  |  | count | At-Lb divergence | divergence.p | count | At-Lb divergence | concordance | divergence.p | concordance.p |
| In(2L)t | 2L | 2225744 | 13154180 | 15 | 0.479968203 | 0.282 | 279394 | 0.426852075 | 0.502912646 | 0.363 | 0.512 |
| In(2R)Ns | 2R | 11278659 | 16163839 | 62 | 0.48336471 | 0.147 | 105545 | 0.427194496 | 0.474853585 | 0.397 | 0.896 |
| In(3L)P | 3L | 3173046 | 16301941 | 3 | 0.242628985 | 0.747 | 290383 | 0.42137602 | 0.502076249 | 0.501 | 0.503 |
| In(3R)C | 3R | 1.60E+07 | 2.60E+07 | 20 | 0.547489936 | 0.08 | 205344 | 0.419489051 | 0.519170927 | 0.56 | 0.182 |
| In(3R)K | 3R | 7576289 | 21966092 | 1 | 0.975097587 | 0.11 | 275032 | 0.417266525 | 0.52398069 | 0.588 | 0.136 |
| In(3R)Mo | 3R | 17232639 | 24857019 | 137 | 0.451377244 | 0.187 | 153064 | 0.415534453 | 0.520780672 | 0.638 | 0.181 |
| In(3R)Payne | 3R | 12257931 | 20569732 | 15 | 0.291547042 | 0.902 | 168293 | 0.419605227 | 0.53104362 | 0.519 | 0.08 |

**Table S3.** List of sites that show divergence between *At* and *Lb* cages in the experiment (effect size >2% and p-value <.05) and that show the most pronounced variation along a cline in eastern North America (clinality FDR<10^−8^). Of these 35 sites, 80% show concordance in the direction of allele change between experimental replicates enriched for AAB and populations that have a higher proportion of AAB in the microbiome.

Table shows: the chromosome arm, position, reference allele, alternate allele, gene name, gene group ID, inversions overlapping the site, coefficient and p-value for divergence between *At* and *Lb* treatments, coefficient and p-value for divergence across clinal populations, concordance between the allele associated with experimental populations and natural populations enriched for the same microbial taxa, average allele frequency in founder, *No-Ad*, *At*, and *Lb*, populations, all annotations of SNP effect according to snpEff, and GO terms. Gene names for SNPs outside genic regions are in parenthesis and indicate the closest gene for gene-counting purposes.

**\*Table is large so it is attached separately.**

**Figure S1:**
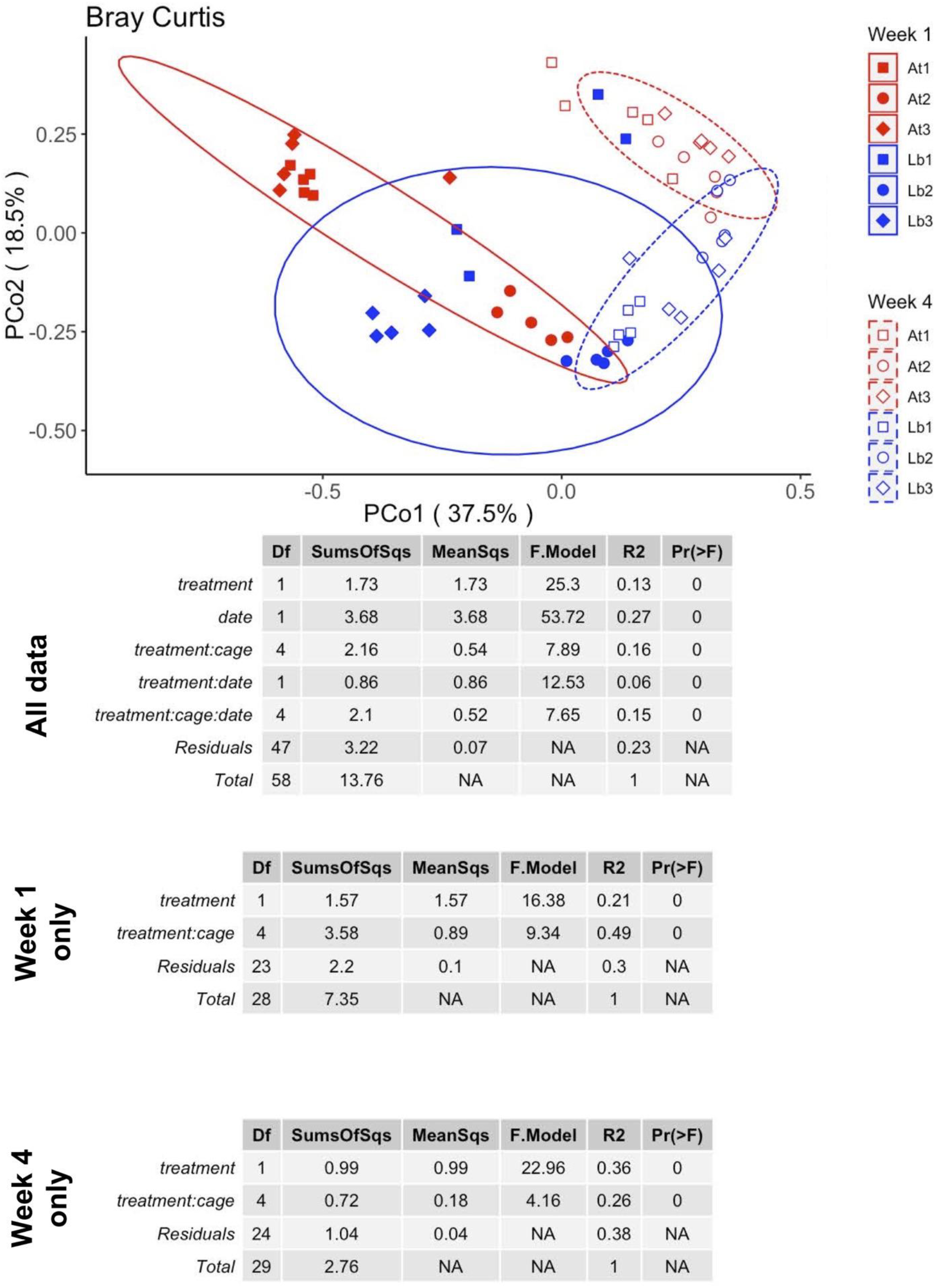
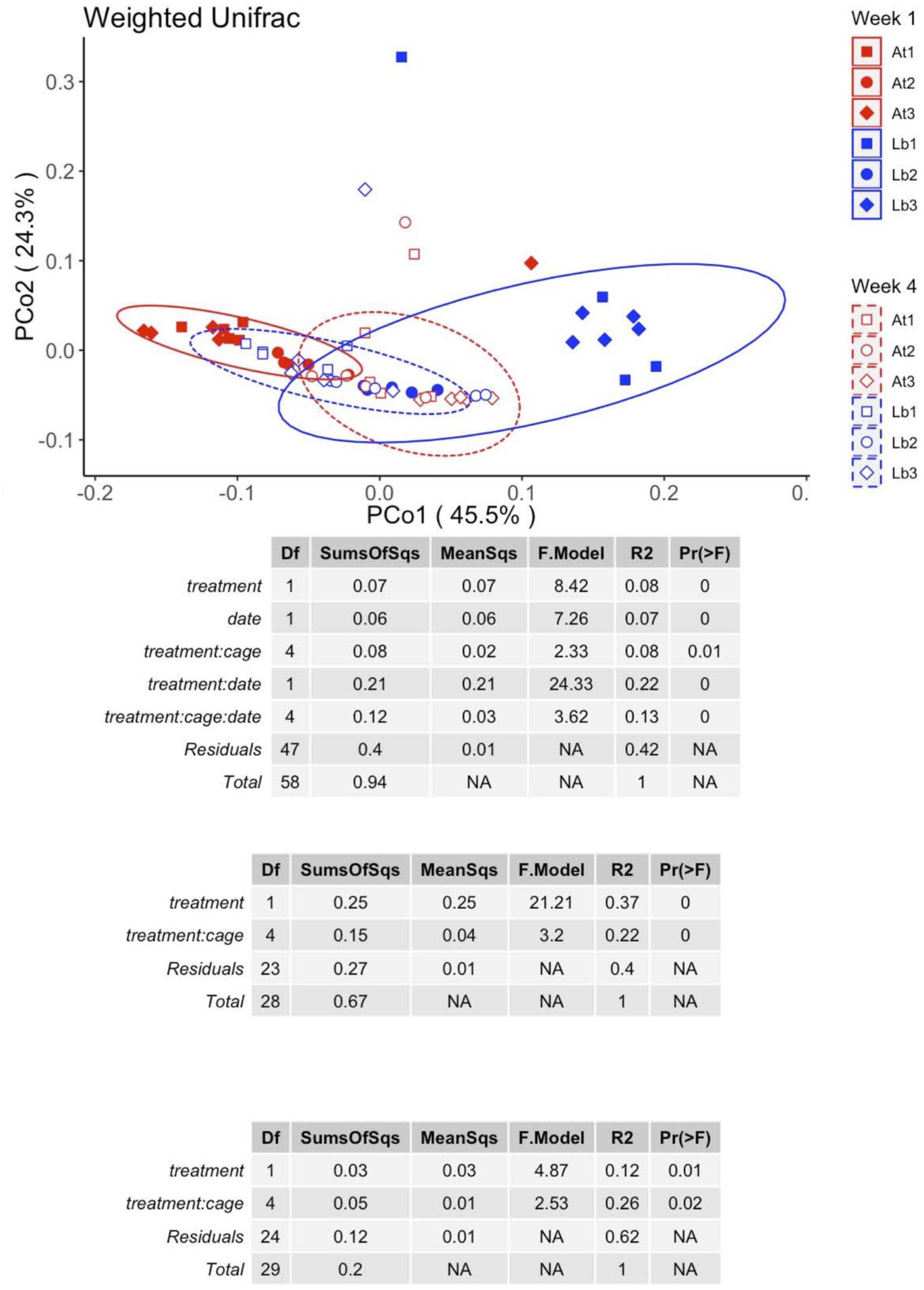
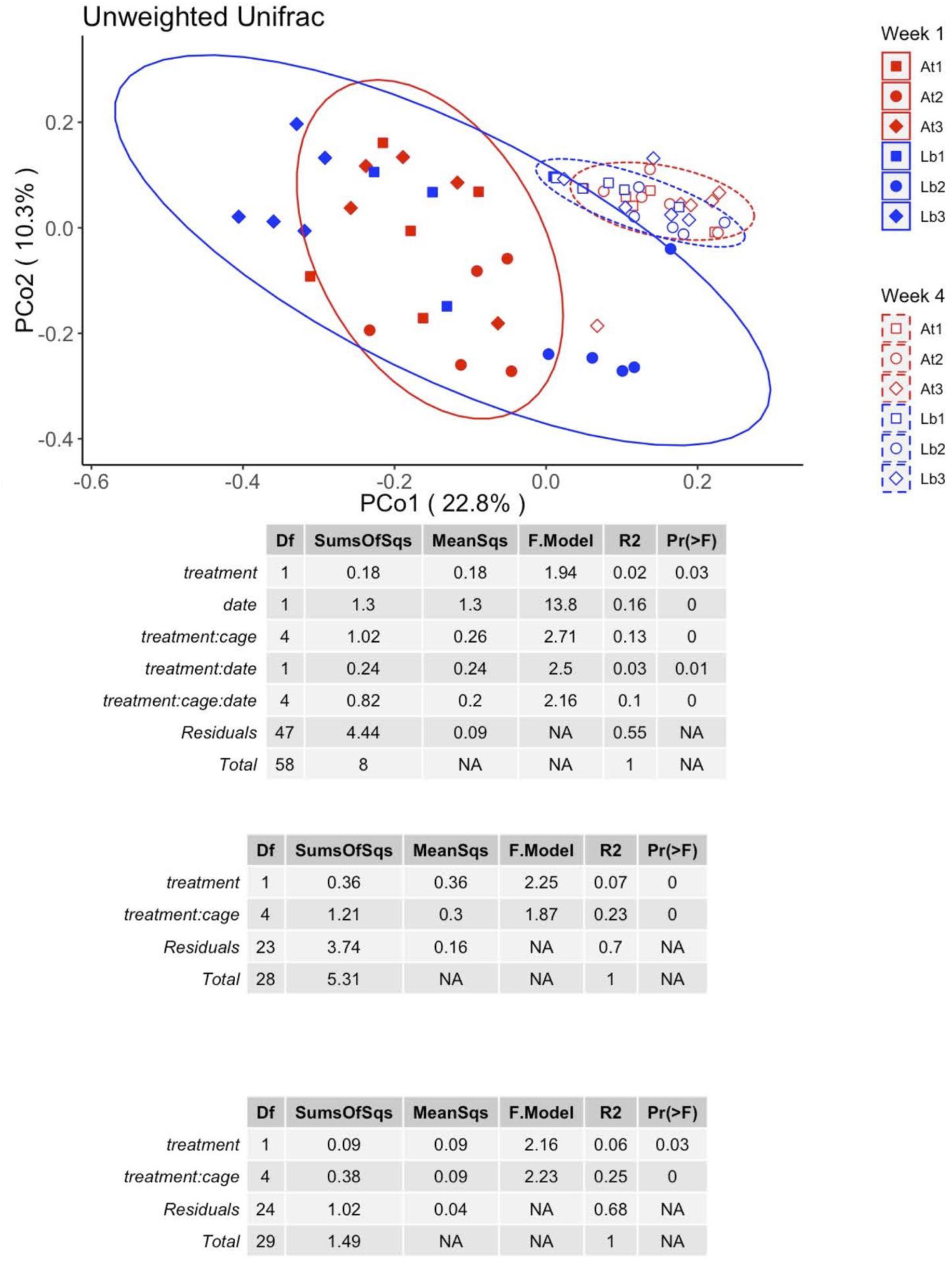
Beta-diversity analysis of microbial communities based on 16S rRNA gene sequencing of whole-body flies collected at week 1 and week 4 of the experiment using Bray-Curtis, weighted Unifrac, and unweighted Unifrac distance metrics. At each time point, five replicate samples of five flies were collected from each cage, the flies were homogenized, and the homogenate was stored at −80°C until a 16S rRNA gene library was prepared. Libraries were sequenced paired-end with 250 bp reads on a Illumina HiSeq. Distance matrices were created using standard QIIME2 parameters. PERMANOVAs were calculated in R using the vegan package.

**Figure S2:**
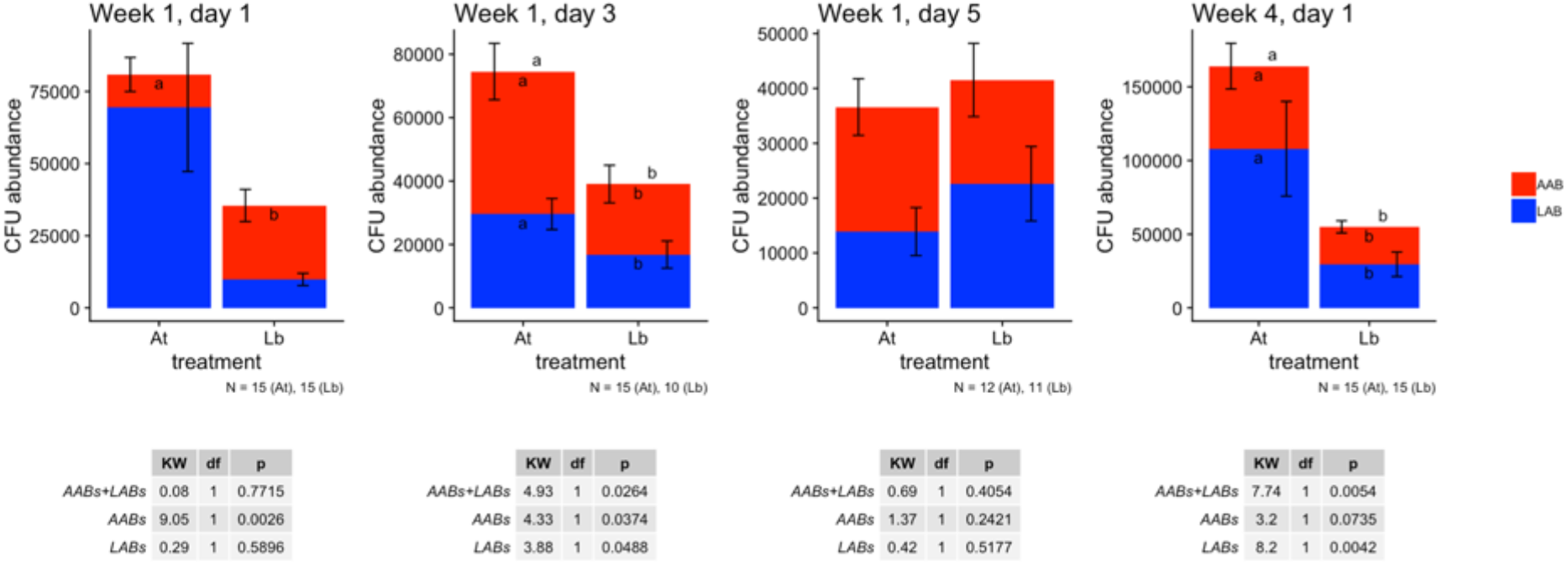
Absolute abundance of CFUs in *D. melanogaster* from outdoor enclosures. Pools of 5 male flies were collected from outdoor enclosures, homogenized in 125 ul phosphate-buffered saline (PBS), and dilution plated on modified de Man-Rogosa-Sharpe medium. Colony forming units (CFUs) were counted on plates after 1-3 days, and normalized to CFU per fly: AAB were copper or tan-colored, while LAB were yellow or white. Values are shown as a mean of all pools, which were collected from each of 3 separate experiments (usually 5 pools per cage per sampling time, exact N shown in caption). For each of three metrics - AAB abundance, LAB abundance, and AAB+LAB abundance - pairwise comparisons between absolute CFU abundances were determined by a Dunn test {dunn.test} and shown by compact letter displays over the bars {rcompanion}. LAB, AAB, and LAB+AAB abundances are shown within the blue bars, within the red bars, and outside of the bars, respectively. If no letters are shown there were no significant differences. The Kruskal-Wallis test statistic (KW), degrees of freedom (df), and p-values (p) for each comparison are reported. All statistics were calculated in R.

**Figure S3:**
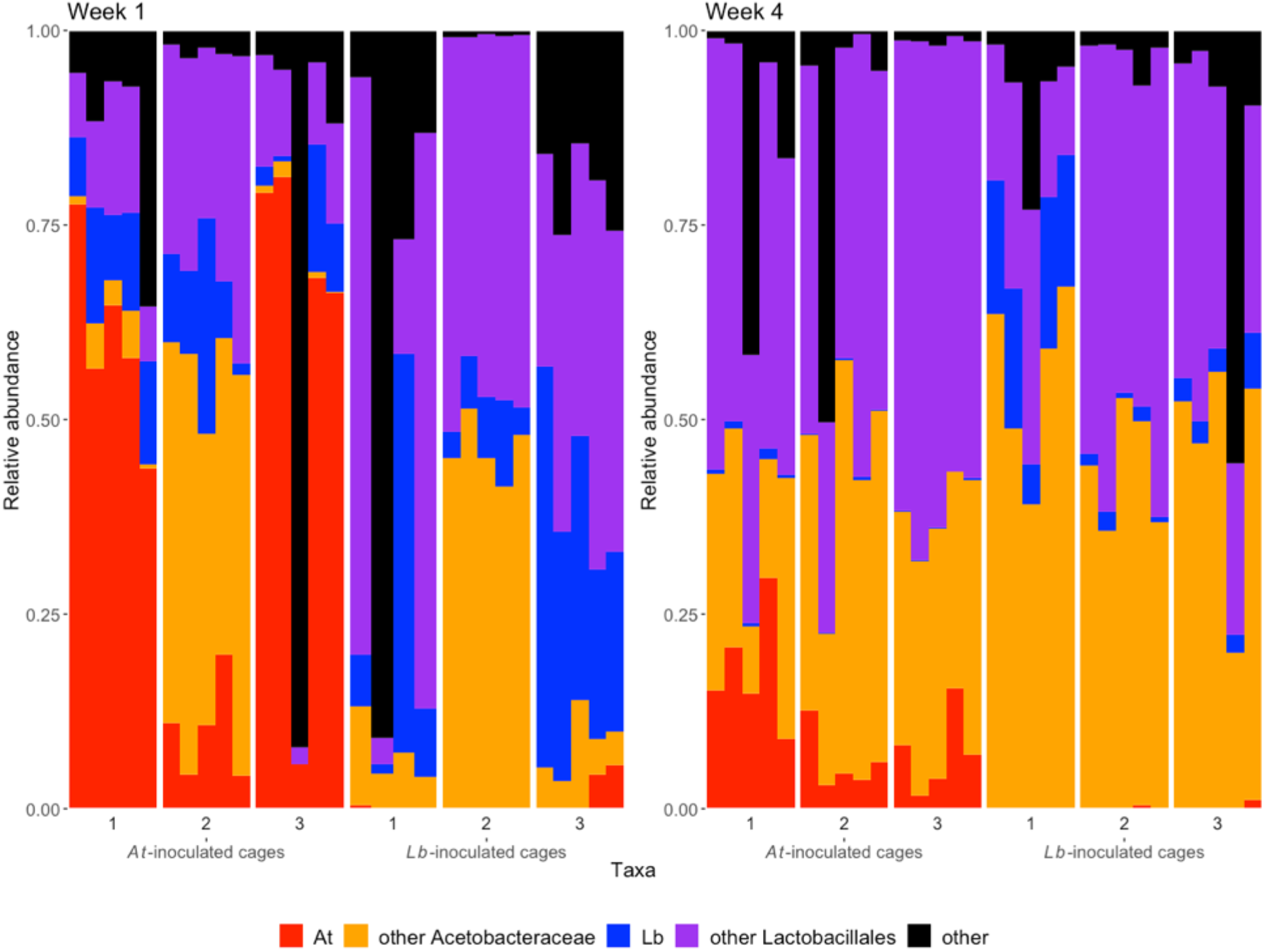
Microbiome composition of homogenized adult flies based on 16S sequence data. Individual bars represent cage-collected pools composed of 5 males with 5 replicate pools sequenced for each cage. Graphs show relative abundance of each microbiome group for each pool (5 from each cage) after 1 week of treatment and 4 weeks of treatment. Flies were sampled from throughout the experimental cages with care taken not to sample flies directly on media.

**Figure S4:**
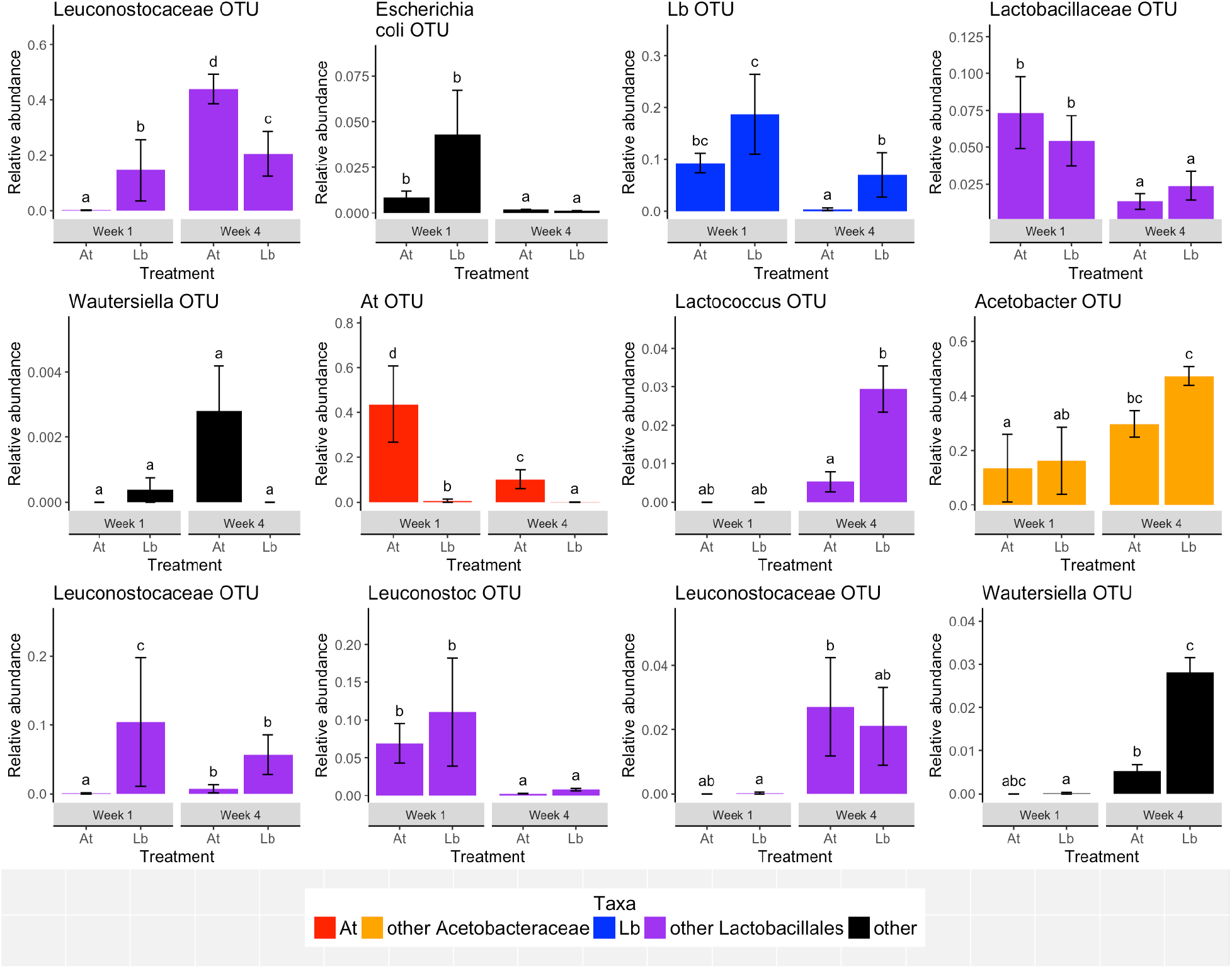
OTUs where we detecting significant differences in abundance between microbiome treatments or within treatments over time as determined by ANCOM with the most stringent correction for multiple tests. Significant differences with treatment and/or time were subsequently confirmed by a mixed effects model with a binomial family and are shown as compact letter displays (different letters represent significant differences between conditions). Most OTUs could not be classified to the species level and are named according to the lowest taxonomic assignment. Each panel shows the relative abundance of a specific OTU as a percentage of total the microbiome community in the *At* and *Lb* experimental treatments at week 1 and week 4 of the experiment. Bars are shaded based on taxonomic assignments and represent the mean +/− SE.

**Figure S5.**
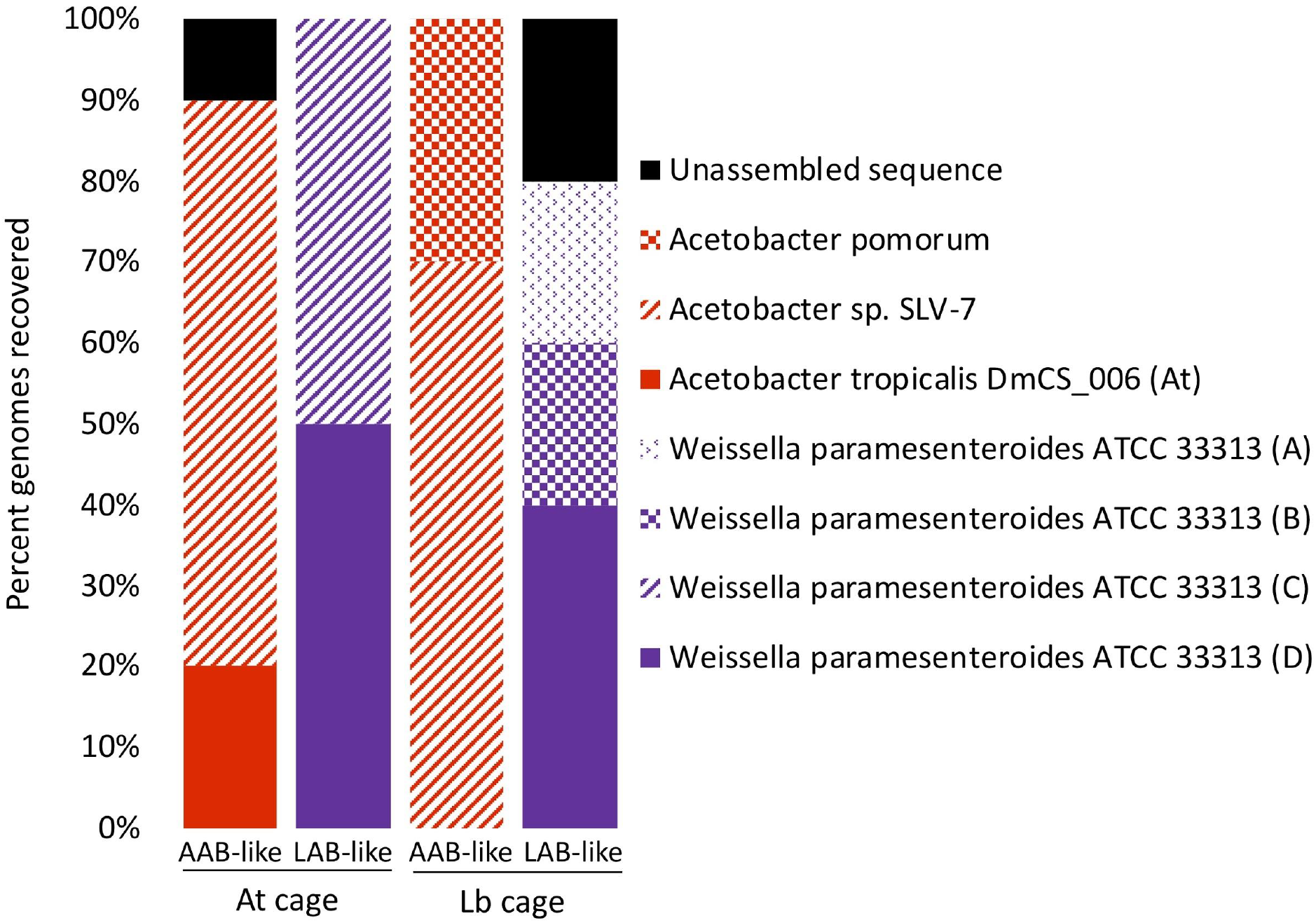
Taxonomic identity of randomly selected bacterial colonies isolated from whole body homogenates of caged *D. melanogaster.* For each of the *At* and *Lb* treatments, 10 AAB-like (copper-colored) and 10 LAB-like (white or yellow colored) colonies were randomly selected, streaked for isolation, and genome sequenced. Genomic libraries were prepared using 800ul of bacteria culture, which was centrifuged at 15,000 × g for 10 minutes. Pellets were resuspended in 600 ul of lysis buffer, and the 600 ul volume was extracted and quantified. In order to fragment the extracted DNA, 1.5 ug of DNA was diluted in 2 ul fragmentase buffer. 2 ul NEB fragmentase was added and each sample was incubated at 37C to digest the fragments to approximately 500 bp (incubation times were optimized to the batch of fragmentase). Enzyme activity was halted after incubation using 10ul (0.25M) EDTA. DNA was cleaned using Zymo DNA Clean and Concentrator 25 columns. For end repair, 50 ul fragmented DNA was mixed with 3 ul NEB enzyme mix, 7ul NEB end prep reaction buffer, and placed in a thermocycler for 30 minutes each for 20C and 65 C. This reaction was combined with 30ul NEB ligation master mix, 1ul ligation enhancer, and 2.5 ul of a unique illumina adapter and then ligated at 20C for 15 minutes. Fragment size selection was done using Ampure SPRI beads (SPB). The bead stock was diluted in a ratio of 109.25 ul SPB to 74.75 ul to ul ddH2O. 160ul diluted SPB were added to 100 ul of end repaired sample, and incubated at 25°C for 5 minutes. The bead-separated supernatant was mixed with 30 ul of SPB stock and the supernatant was discarded. Samples were washed twice with 200ul 80% EtOH, resuspended in 22.5 ul RSB for 5 min, and 17.5 ul was transferred to an ALP plate. Barcode ligation was enriched with the KAPA Library Amplification kit according to manufacturer instructions, size selection was repeated, libraries normalized to 5ng/uL via Qubit, and sequenced by 2×125 bp sequencing on a Illumina HiSeq 2500 at the BYU DNA sequencing center. Genome sequences were assembled using Velvet 1.2.10 as described previously (Newell, 2014). Briefly, the nucleotide coverage of each of the raw assemblies was determined based on the size of the expected genome (A. tropicalis DmCs_006 or L. brevis DmCs_002), assembled into contigs across a kmer range of 85-123, adjusted for expected coverage and coverage cutoff, and a single assembly that minimized node number while maximizing n50 value and total genome size was selected manually. When sequence reads provided greater than 200X nucleotide coverage, the raw reads were split evenly into subsets providing 101-200X coverage, and each subset was assembled as described above except that there were no adjustments for expected coverage and coverage cutoff. The contigs file from the multiple sub-assemblies for each genome were used to create a final assembly. The similarity of each of these genomes was compared in an all against all mummer comparison that included the reference genomes for *A. tropicalis* DmCs_006 or *L. brevis* DmCs_002. Genomes were clustered according to a 99.9% or greater M-to-M score. Taxonomic identity of the representative sequence for each cluster, picked as the assembly with the fewest contigs, was assigned using JSpecies. **Taxonomic identities of bacterial isolates recovered from evolving flies.** A pool of five male flies from a randomly-selected cage that had been inoculated with either *At* or *Lb* was collected, homogenized, and plated on mMRS. From each pool, ten AAB-like colonies and ten LAB-like colonies were cultured in isolation and subjected to whole genome sequencing. Whole genome nucleotide similarity of the strains was determined by ANIm, with different isolates assigned at ANIM>99.9%. The results show that strains identical to the inoculated *At* strain could be recovered from the treatment flies, confirming that the treatment bacteria were colonizing the flies.

**Fig S6:**
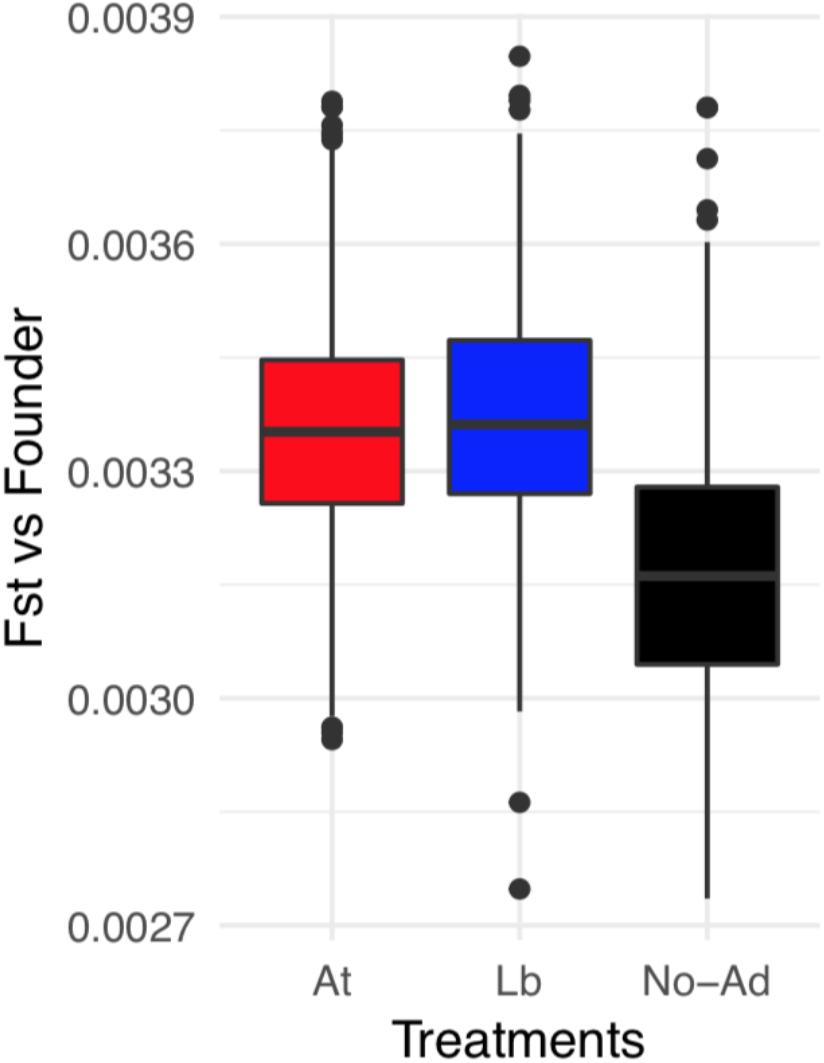
Average Fst between the founder population and each treatment for 1,000 subsets of 1,000 segregating sites. For all comparisons, allele frequencies at each site were averaged within replicate populations of the same treatment before calculating Fst. For the *No-Ad* treatment, a random sample of 3 replicate *No-Ad* populations was chosen for each site subset to match the number of *At* and *Lb* replicates.

**Figure S7:**
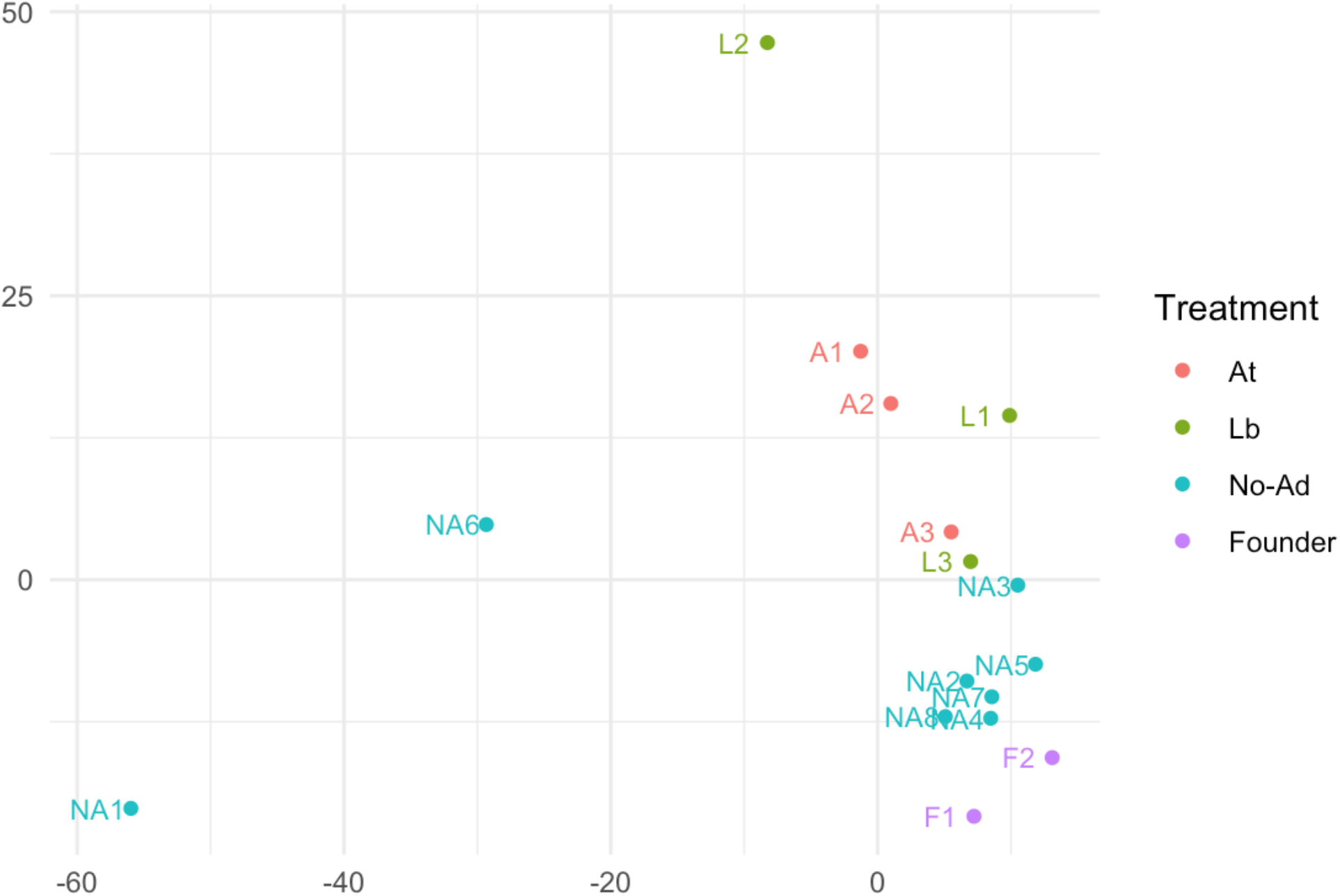
PCA based on all SNPs for all experimental cages and the founding population colored by treatment. ‘Founder’ represent sub-samples of the initial population used to found each replicate and were sampled at Day 0. All outdoor treatment cages were sampled at Day 45.

