## Supplemental Table 1 for "Microbiome composition shapes rapid genomic adaptation of *Drosophila melanogaster*"

| chrm | pos | ref | alt | gene | p.At-Lb_div | afMean.Founder | afMean.NoAd | afMean.At | afMean.Lb | p.clinality | GOterm |
|---|---|---|---|---|---|---|---|---|---|---|---|
| 2L | 409409 | A | G | alpha-Adaptin | 4.65E-06 | 0.222 | 0.281 | 0.306 | 0.219 | 2.58E-01 | protein transporter activity;clathrin adaptor activity;endocytosis;mitotic cleavage furrow ingression;ovarian follicale cell development;positive regulation of autophagy |
| 2L | 469463 | C | A | (MED15) | 7.30E-05 | 0.124 | 0.142 | 0.121 | 0.068 | 1.11E-01 | NA |
| 2L | 1101026 | C | G | CG4629 | 1.40E-05 | 0.442 | 0.416 | 0.285 | 0.471 | 6.94E-01 | protein serine/threonine kinase activity;ATP binding;protein phosphorylation;cell adhesion;regulation of cell shape |
| 2L | 1674858 | C | T | chinmo | 5.48E-05 | 0.228 | 0.191 | 0.151 | 0.245 | 2.25E-01 | DNA-binding transcription factor activity;imaginal disc-derived wing morphogenesis;mushroom body development;metal ion binding;dendrite morphogenesis |
| 2L | 1998051 | T | A | CG33543 | 1.61E-05 | 0.248 | 0.233 | 0.257 | 0.220 | 8.28E-05 | cell adhesion;membrane |
| 2L | 2087653 | A | G | dpr3 | 6.96E-05 | 0.412 | 0.434 | 0.463 | 0.299 | 1.67E-01 | store-operated calcium entry;sensory perception of chemical stimulus |
| 2L | 2334084 | A | G | CG9967 | 2.69E-05 | 0.270 | 0.225 | 0.184 | 0.276 | 7.93E-01 | |
| 2L | 2669544 | T | A | (Gr23a) | 4.15E-05 | 0.110 | 0.074 | 0.057 | 0.161 | 1.84E-01 | NA |
| 2L | 3433591 | T | G | (pgant2) | 6.77E-05 | 0.088 | 0.100 | 0.076 | 0.136 | 5.67E-01 | NA |
| 2L | 3852967 | A | C | (slp2) | 9.60E-05 | 0.563 | 0.543 | 0.632 | 0.478 | 6.96E-01 | NA |
| 2L | 4132809 | A | T | (Sr) | 1.50E-04 | 0.043 | 0.063 | 0.034 | 0.091 | 9.99E-01 | NA |
| 2L | 4313991 | T | A | tutl | 1.15E-04 | 0.020 | 0.057 | 0.025 | 0.046 | 6.49E-01 | axon guidance;axonal defasciculation;peripheral nervous system development;flight behavior;mechanosensory behavior;synaptic target recognition;adult locomotory behavior;axon midline choice point recognition;larval behavior;lateral inhibition;regulation of dendrite morphogenesis;dendrite self-avoidance |
| 2L | 4814724 | T | A | CG15629 | 8.35E-05 | 0.116 | 0.119 | 0.150 | 0.080 | 3.63E-01 | metabolic process;oxidoreductase activity |

| | | | | | | | | | | | |
|---|---|---|---|---|---|---|---|---|---|---|---|
| 2L | 4867676 | A | G | pog | 4.54E-06 | 0.212 | 0.269 | 0.205 | 0.300 | 1.92E-01 | regulation of antimicrobial peptide biosynthetic process;G protein-coupled receptor activity;G protein-coupled receptor signaling pathway;G protein-coupled glutamate receptor signaling pathway;germ-band extension;glutamate receptor activity;integral component of membrane |
| 2L | 5347750 | G | C | nompC | 1.24E-04 | 0.076 | 0.068 | 0.091 | 0.063 | 6.43E-01 | startle response;ion channel activity;cation channel activity;calcium channel activity;cilium;cation transport;calcium ion transport;sensory perception of sound;mechanosensory behavior;cytoskeletal protein binding;mechanosensitive ion channel activity;response to auditory stimulus;dendrite;ankyrin binding;cation channel complex;locomotion;neuronal cell body;sensory perception of mechanical stimulus;detection of mechanical stimulus involved in sensory perception;sensory perception of touch;calcium ion transmembrane transport |
| 2L | 5870688 | A | G | CG8965 | 2.21E-06 | 0.075 | 0.072 | 0.025 | 0.080 | 5.65E-02 | protein homodimerization activity; Ras GTPase binding; glial cell development; prisitive regulation of epidermal growth factor receptor signaling pathway R7 cell development |
| 2L | 6256166 | G | A | CG9486 | 1.49E-04 | 0.110 | 0.078 | 0.095 | 0.124 | 4.52E-01 | N-acetyltransferase activity |
| 2L | 6499556 | T | C | CG31637 | 8.75E-05 | 0.754 | 0.742 | 0.714 | 0.814 | 7.04E-04 | sulfotransferase activity |
| 2L | 6603495 | G | A | (CG11319) | 9.78E-05 | 0.350 | 0.355 | 0.295 | 0.432 | NA | NA |
| 2L | 7392478 | T | C | CG5160 | 1.91E-05 | 0.063 | 0.106 | 0.056 | 0.106 | 2.62E-02 | GTPase activity;GTP binding;obsolete GTP catabolic process;small GTPase mediated signal transduction;membrane;neurogenesis |

| | | | | | | | | | | | |
|---|---|---|---|---|---|---|---|---|---|---|---|
| 2L | 7417601 | G | A | chm | 1.40E-05 | 0.185 | 0.181 | 0.220 | 0.130 | 6.82E-01 | DNA-binding transcription factor activity;transcription coactivator activity;histone acetyltransferase activity;nucleus;transcription factor binding;zinc ion binding;negative regulation of cell fate specification;H4 histone acetyltransferase activity;sensory organ precursor cell fate determination;gene silencing;chaeta development;negative regulation of transcription |
| 2L | 7435458 | T | A | (CG5958) | 1.26E-04 | 0.116 | 0.105 | 0.162 | 0.097 | 2.19E-01 | NA |
| 2L | 7462699 | G | A | (Gr28b) | 1.05E-05 | 0.052 | 0.047 | 0.081 | 0.048 | 4.20E-01 | NA |
| 2L | 7638255 | G | T | (Slob) | 1.14E-04 | 0.140 | 0.158 | 0.116 | 0.207 | NA | NA |
| 2L | 8211227 | A | T | (Ssb-c31a) | 7.34E-06 | 0.035 | 0.016 | 0.016 | 0.042 | NA | NA |
| 2L | 8304548 | A | C | Scgalpha | 7.24E-05 | 0.038 | 0.037 | 0.027 | 0.059 | 9.35E-01 | calcium ion binding;structural constituent of muscle;sarcoglycan complex |
| 2L | 8784561 | T | C | (CG9468) | 3.68E-05 | 0.875 | 0.879 | 0.920 | 0.803 | 3.28E-01 | NA |
| 2L | 9102902 | G | A | (CG31609) | 8.59E-05 | 0.196 | 0.209 | 0.156 | 0.271 | 6.33E-01 | NA |
| 2L | 9185493 | A | T | tai | 1.10E-04 | 0.017 | 0.008 | 0.032 | 0.011 | 1.00E+00 | transcription coactivator activity;obsolete signal transducer activity;nucleus;regulation of transcription |
| 2L | 9254319 | A | G | lectin-30A | 2.33E-05 | 0.333 | 0.250 | 0.357 | 0.195 | 4.60E-01 | galactose binding;extracellular space;carbohydrate binding;multicellular organism reproduction |
| 2L | 9365904 | C | T | (CG34181) | 1.51E-04 | 0.056 | 0.095 | 0.043 | 0.132 | 5.87E-01 | NA |

| | | | | | | | | | | | |
|---|---|---|---|---|---|---|---|---|---|---|---|
| 2L | 9447685 | C | T | numb | 1.21E-04 | 0.119 | 0.150 | 0.083 | 0.167 | 9.94E-01 | cell fate determination;nucleic acid binding;Notch binding;protein binding;nucleus;cytoplasm;cell cortex;Notch signaling pathway;neuroblast fate determination;neuroblast proliferation;central nervous system development;ventral cord development;peripheral nervous system development;heart development;rhythmic behavior;protein localization;glial cell migration;asymmetric cell division;regulation of Notch signaling pathway;regulation of asymmetric cell division;sensory organ precursor cell fate determination;embryonic heart tube development;muscle cell fate specification;sensory organ precursor cell division;cell fate commitment;basal part of cell;basal cortex;negative regulation of Notch signaling pathway;positive regulation of endocytosis;regulation of neurogenesis;centrosome localization;regulation of nervous system development;asymmetric neuroblast division;pericardial nephrocyte differentiation;Malpighian tubule tip cell differentiation |
| 2L | 9772922 | G | C | ppk16 | 7.39E-05 | 0.049 | 0.040 | 0.024 | 0.067 | 2.30E-01 | sodium channel activity;sodium ion transport;integral component of membrane |
| 2L | 9783859 | G | C | IP3K1 | 1.19E-04 | 0.173 | 0.139 | 0.118 | 0.185 | 3.62E-01 | calmodulin binding;cytoplasm;response to oxidative stress;inositol-1 |
| 2L | 9948936 | G | T | (CG5853) | 6.81E-05 | 0.105 | 0.139 | 0.095 | 0.223 | 3.47E-02 | NA |
| 2L | 1E+07 | A | G | CG42843 | 1.22E-04 | 0.143 | 0.169 | 0.100 | 0.227 | 4.17E-01 | cell-cell adhesion mediator activity; G protein-coupled receptor activity; axon guidance; dendrite self-avoidance; homophilic cell adhesion |

| | | | | | | | | | | | |
|---|---|---|---|---|---|---|---|---|---|---|---|
| 2L | 1E+07 | G | A | CG5355 | 2.90E-06 | 0.041 | 0.035 | 0.029 | 0.070 | 7.65E-01 | serine-type endopeptidase activity;proteolysis;carboxylic ester hydrolase activity;serine-type exopeptidase activity |
| 2L | 1.2E+07 | A | T | (CG31706) | 2.85E-05 | 0.099 | 0.122 | 0.144 | 0.077 | NA | NA |
| 2L | 1.2E+07 | T | A | Pde1c | 7.77E-05 | 0.242 | 0.206 | 0.237 | 0.176 | 8.84E-01 | calcium- and calmodulin-regulated 3',5'-cyclic-AMP/GMP phosphodiesterase activity; calmodulin binding; male mating behavior; signal transduction |
| 2L | 1.2E+07 | G | A | bru-2 | 1.25E-04 | 0.040 | 0.054 | 0.072 | 0.010 | 1.10E-01 | nucleotide binding;RNA binding;mRNA binding;negative regulation of translation |
| 2L | 1.3E+07 | A | T | (bun) | 5.72E-05 | 0.284 | 0.285 | 0.182 | 0.242 | 2.34E-01 | NA |
| 2L | 1.3E+07 | C | A | CG15484 | 1.29E-04 | 0.013 | 0.031 | 0.049 | 0.012 | 4.73E-01 | |
| 2L | 1.3E+07 | A | T | (CG15483) | 1.95E-05 | 0.058 | 0.086 | 0.093 | 0.048 | 4.35E-01 | NA |
| 2L | 1.3E+07 | C | A | (kek1) | 6.80E-05 | 0.118 | 0.141 | 0.195 | 0.090 | 3.85E-01 | NA |
| 2L | 1.3E+07 | A | C | CG42784 | 3.30E-05 | 0.207 | 0.189 | 0.179 | 0.294 | 1.93E-01 | |
| 2L | 1.4E+07 | C | G | (nimB4) | 1.54E-05 | 0.070 | 0.059 | 0.071 | 0.026 | 8.58E-01 | NA |
| 2L | 1.4E+07 | G | A | rk | 4.75E-05 | 0.634 | 0.705 | 0.599 | 0.757 | 9.03E-01 | epithelial to mesenchymal transition;G protein-coupled receptor activity;plasma membrane;integral component of plasma membrane;G protein-coupled receptor signaling pathway;neuropeptide signaling pathway;regulation of chitin-based cuticle tanning;neuropeptide receptor activity;integral component of membrane;protein-hormone receptor activity |
| 2L | 1.4E+07 | G | T | ıAcRalpha-34I | 7.24E-05 | 0.040 | 0.037 | 0.012 | 0.054 | 4.39E-01 | acethylcholine-gated cation-selective channel activity;transmembrane signaling receptor;cation transport;chemical synaptic transmission;ion transmembrane transport;nervous system process;regulation of membrane potential |

| | | | | | | | | | | | |
|---|---|---|---|---|---|---|---|---|---|---|---|
| 2L | 1.5E+07 | C | T | GABA-B-R1 | 1.39E-04 | 0.158 | 0.231 | 0.158 | 0.253 | 5.37E-01 | G protein-coupled GABA receptor activity;integral component of plasma membrane;G protein-coupled receptor signaling pathway;negative regulation of adenylate cyclase activity;gamma-aminobutyric acid signaling pathway;integral component of membrane;G protein-coupled receptor heterodimeric complex;protein heterodimerization activity |
| 2L | 1.6E+07 | A | T | (l_2_35Di) | 9.44E-05 | 0.153 | 0.178 | 0.214 | 0.128 | 2.75E-01 | NA |
| 2L | 1.6E+07 | G | A | (CG7653) | 2.51E-05 | 0.147 | 0.159 | 0.178 | 0.107 | 1.66E-01 | NA |
| 2L | 1.6E+07 | G | A | (beat-Ia) | 1.51E-04 | 0.028 | 0.053 | 0.022 | 0.050 | 7.62E-01 | NA |
| 2L | 1.6E+07 | G | C | (CG34168) | 1.33E-04 | 0.141 | 0.145 | 0.187 | 0.093 | 6.22E-01 | NA |
| 2L | 1.6E+07 | C | T | Ca-alpha1D | 5.70E-06 | 0.170 | 0.159 | 0.251 | 0.122 | 2.63E-01 | voltage-gated calcium channel activity;voltage-gated calcium channel complex;calcium ion transport;muscle contraction;embryo development;basolateral plasma membrane;apical plasma membrane;epithelial fluid transport;calcium ion transmembrane transport |
| 2L | 1.6E+07 | G | A | CG42818 | 6.96E-05 | 0.066 | 0.093 | 0.118 | 0.078 | 3.78E-02 | |
| 2L | 1.7E+07 | G | C | CG42389 | 7.61E-05 | 0.262 | 0.212 | 0.164 | 0.240 | 7.62E-01 | |
| 2L | 1.7E+07 | G | A | trpgamma | 4.10E-06 | 0.640 | 0.673 | 0.599 | 0.668 | 1.67E-03 | cation channel activity;mechanosensitive ion channel activity;calcium channel activity;inositol 1,4,5 triphosphate binding;adult walking behavior;neuromuscular process controlling posture |
| 2L | 1.8E+07 | T | C | (CR43304) | 1.45E-04 | 0.062 | 0.110 | 0.091 | 0.147 | NA | NA |

| | | | | | | | | | | | |
|---|---|---|---|---|---|---|---|---|---|---|---|
| 2L | 1.8E+07 | A | C | CadN2 | 1.27E-07 | 0.279 | 0.321 | 0.370 | 0.199 | 6.71E-01 | NA;calcium ion binding;plasma membrane;integral component of plasma membrane;homophilic cell adhesion via plasma membrane adhesion molecules;ommatidial rotation;calcium-dependent cell-cell adhesion via plasma membrane cell adhesion molecules;R8 cell development;R7 cell development;axon extension involved in axon guidance;cell adhesion molecule binding |
| 2L | 1.8E+07 | C | T | (CadN2) | 9.05E-05 | 0.114 | 0.070 | 0.042 | 0.107 | 4.11E-02 | NA |
| 2L | 1.8E+07 | T | C | rdo | 1.09E-04 | 0.313 | 0.355 | 0.480 | 0.387 | 3.07E-01 | ocellus development |
| 2L | 1.8E+07 | A | G | CG42750 | 4.44E-05 | 0.013 | 0.032 | 0.027 | 0.078 | 3.08E-01 | proteolysis;metalloexopeptidase activity;dipeptidyl-peptidase activity;dipeptidase activity |
| 2L | 1.8E+07 | C | T | (CG31787) | 7.40E-06 | 0.481 | 0.394 | 0.326 | 0.526 | 8.20E-04 | NA |
| 2L | 1.8E+07 | C | T | (CG31787) | 6.37E-06 | 0.490 | 0.398 | 0.321 | 0.536 | 7.18E-04 | NA |
| 2L | 1.8E+07 | A | G | (CG31787) | 1.33E-04 | 0.023 | 0.023 | 0.036 | 0.010 | 7.82E-01 | NA |
| 2L | 1.9E+07 | A | G | Pde11 | 8.90E-05 | 0.483 | 0.464 | 0.410 | 0.508 | 2.33E-01 | 3',5'-cyclic-AMP/GMP phosphodiesterase activity |
| 2L | 1.9E+07 | T | C | (tup) | 1.08E-04 | 0.855 | 0.860 | 0.898 | 0.820 | 3.48E-01 | NA |
| 2L | 1.9E+07 | T | A | CG10650 | 5.75E-05 | 0.039 | 0.038 | 0.029 | 0.066 | 1.00E+00 | |
| 2L | 1.9E+07 | G | A | Ddc | 6.03E-05 | 0.512 | 0.534 | 0.512 | 0.587 | 7.52E-02 | aromatic-L-amino-acid decarboxylase activity;catecholamine metabolic process;dopamine biosynthetic process from tyrosine;serotonin biosynthetic process from tryptophan;learning or memory;anesthesia-resistant memory;long-term memory;courtship behavior;eclosion rhythm;response to wounding;pyridoxal phosphate binding;wing disc development;growth;thermosensory behavior;thermotaxis;developmental pigmentation;regulation of adult chitin-containing cuticle pigmentation;adult chitin-containing cuticle pigmentation |

| | | | | | | | | | | | |
|---|---|---|---|---|---|---|---|---|---|---|---|
| 2L | 1.9E+07 | T | C | drl | 4.68E-06 | 0.066 | 0.067 | 0.094 | 0.028 | 3.62E-02 | protein tyrosine kinase activity;transmembrane receptor protein tyrosine kinase activity;ATP binding;plasma membrane;protein phosphorylation;signal transduction;axon guidance;salivary gland morphogenesis;haltere development;learning or memory;memory;olfactory learning;axon midline choice point recognition;muscle attachment;determination of muscle attachment site;Wnt-protein binding;axon |
| 2L | 1.9E+07 | C | G | drl | 4.45E-05 | 0.349 | 0.362 | 0.419 | 0.357 | 8.97E-01 | protein tyrosine kinase activity;transmembrane receptor protein tyrosine kinase activity;ATP binding;plasma membrane;protein phosphorylation;signal transduction;axon guidance;salivary gland morphogenesis;haltere development;learning or memory;memory;olfactory learning;axon midline choice point recognition;muscle attachment;determination of muscle attachment site;Wnt-protein binding;axon |
| 2L | 2E+07 | C | T | bsh | 1.04E-04 | 0.078 | 0.095 | 0.133 | 0.084 | 8.22E-04 | transcription regulatory region sequence-specific DNA binding;DNA-binding transcription factor activity;nucleus;regulation of transcription |
| 2L | 2.1E+07 | A | T | (Oseg5) | 1.39E-04 | 0.248 | 0.255 | 0.360 | 0.246 | 8.32E-01 | NA |
| 2L | 2.2E+07 | C | A | (tsh) | 4.04E-05 | 0.050 | 0.037 | 0.046 | 0.122 | NA | NA |
| 2R | 1412432 | A | G | CG30438 | 1.08E-04 | 0.017 | 0.063 | 0.053 | 0.033 | 5.97E-03 | metabolic process;transferase activity |
| 2R | 2476210 | T | C | jing | 3.32E-05 | 0.095 | 0.085 | 0.068 | 0.147 | 3.73E-01 | negative regulation of transcription by RNA polymerase II;nucleic acid binding;DNA-binding transcription factor activity;nucleus;regulation of transcription |
| 2R | 3527213 | T | C | CG43340 | 2.11E-05 | 0.743 | 0.808 | 0.845 | 0.792 | 9.22E-01 | intracellular protein transport;imaginal disc-derived wing morphogenesis;Rab GTPase binding |
| 2R | 4166659 | T | C | (CG30371) | 5.47E-05 | 0.094 | 0.063 | 0.095 | 0.049 | 1.10E-02 | NA |

| | | | | | | | | | | | |
|---|---|---|---|---|---|---|---|---|---|---|---|
| 2R | 4180261 | A | C | (CG30371) | 7.55E-05 | 0.895 | 0.906 | 0.929 | 0.852 | 4.12E-01 | NA |
| 2R | 4237786 | A | G | pdm3 | 1.49E-04 | 0.027 | 0.028 | 0.019 | 0.062 | 5.67E-01 | DNA-binding transcription factor activity;nucleus;regulation of transcription |
| 2R | 4684621 | A | G | (sns) | 3.90E-05 | 0.024 | 0.033 | 0.011 | 0.047 | 8.97E-02 | NA |
| 2R | 5046734 | G | C | CG8008 | 1.23E-04 | 0.554 | 0.473 | 0.440 | 0.570 | 2.02E-01 | integral component of membrane;transmembrane transport |
| 2R | 5486967 | C | T | clos | 4.55E-05 | 0.334 | 0.377 | 0.320 | 0.406 | 4.07E-03 | extracellular region;vitelline membrane formation involved in chorion-containing eggshell formation;terminal region determination;torso signaling pathway |
| 2R | 5941384 | T | C | magu | 1.13E-04 | 0.417 | 0.456 | 0.403 | 0.587 | 5.12E-04 | calcium ion binding;extracellular region;NA;determination of adult lifespan;external side of plasma membrane;regulation of BMP signaling pathway;positive regulation of BMP signaling pathway;germ-line stem cell population maintenance;heparan sulfate proteoglycan binding;regulation of imaginal disc-derived wing size |
| 2R | 6289071 | A | C | CG42732 | 1.10E-04 | 0.143 | 0.154 | 0.143 | 0.241 | 2.11E-01 | potassium channel activity;potassium ion transport;calcium-activated potassium channel activity;membrane;neurogenesis;neuron projection morphogenesis |
| 2R | 6321928 | C | T | RanBPM | 1.13E-05 | 0.091 | 0.116 | 0.092 | 0.129 | 6.93E-02 | cytoplasm;cytoskeleton organization;receptor signaling pathway via JAK-STAT;ovarian follicle cell-cell adhesion;Ran GTPase binding;larval feeding behavior;dorsal appendage formation;germ-line stem-cell niche homeostasis |

| | | | | | | | | | | | |
|---|---|---|---|---|---|---|---|---|---|---|---|
| 2R | 6322353 | T | C | RanBPM | 8.88E-05 | 0.131 | 0.138 | 0.144 | 0.113 | 8.76E-01 | cytoplasm;cytoskeleton organization;receptor signaling pathway via JAK-STAT;ovarian follicle cell-cell adhesion;Ran GTPase binding;larval feeding behavior;dorsal appendage formation;germ-line stem-cell niche homeostasis |
| 2R | 6384019 | T | G | lola | 1.40E-04 | 0.807 | 0.737 | 0.770 | 0.650 | 6.70E-01 | startle response;inter-male aggressive behavior;nucleic acid binding;DNA-binding transcription factor activity;protein binding;nucleus;regulation of transcription |
| 2R | 6679768 | A | G | mthl13 | 1.21E-04 | 0.896 | 0.914 | 0.879 | 0.955 | 7.71E-02 | G protein-coupled receptor activity;G protein-coupled receptor signaling pathway;integral component of membrane |
| 2R | 6746358 | T | G | CG30015 | 1.41E-04 | 0.728 | 0.701 | 0.669 | 0.808 | 2.95E-01 | |
| 2R | 7545689 | A | G | Roc2 | 1.15E-05 | 0.195 | 0.200 | 0.241 | 0.155 | 7.51E-01 | ubiquitin-protein transferase activity;zinc ion binding;protein ubiquitination |
| 2R | 7630421 | T | C | (pyr) | 1.43E-04 | 0.045 | 0.043 | 0.023 | 0.048 | 3.86E-01 | NA |
| 2R | 7669315 | G | A | ths | 6.59E-05 | 0.091 | 0.115 | 0.153 | 0.095 | 4.22E-02 | fibroblast growth factor receptor binding;hindgut morphogenesis;mesoderm development;heart development;larval visceral muscle development;larval somatic muscle development;growth factor activity;fibroblast growth factor receptor signaling pathway;glial cell differentiation;myoblast migration |
| 2R | 8090152 | G | C | (CG8858) | 1.07E-04 | 0.087 | 0.107 | 0.073 | 0.133 | 5.36E-01 | NA |
| 2R | 8300037 | T | C | (Cpr49Ah) | 2.92E-05 | 0.606 | 0.638 | 0.702 | 0.617 | 6.15E-02 | NA |

| | | | | | | | | | | | |
|---|---|---|---|---|---|---|---|---|---|---|---|
| 2R | 8669385 | T | C | sca | 1.12E-04 | 0.023 | 0.036 | 0.024 | 0.065 | 1.81E-01 | obsolete signal transducer activity;extracellular region;fibrinogen complex;nervous system development;R8 cell fate commitment;chaeta morphogenesis;imaginal disc-derived wing margin morphogenesis;ommatidial rotation;female meiosis chromosome segregation;regulation of R8 cell spacing in compound eye;lateral inhibition;compound eye development;response to alcohol |
| 2R | 9412636 | G | A | Vmat | 2.69E-06 | 0.073 | 0.104 | 0.201 | 0.105 | 7.43E-01 | obsolete synaptic vesicle amine transmembrane transporter activity;neurotransmitter transport;synaptic vesicle;monoamine transmembrane transporter activity;drug transmembrane transporter activity;aminergic neurotransmitter loading into synaptic vesicle;monoamine transport;dopamine transport;integral component of membrane;neuromuscular junction;histamine transport;transmembrane transport |
| 2R | 9609003 | T | A | fas | 1.49E-04 | 0.072 | 0.091 | 0.056 | 0.134 | NA | morphogenesis of an epithelium;dorsal closure;salivary gland development;salivary gland morphogenesis;foregut morphogenesis;Malpighian tubule morphogenesis;cardioblast cell fate determination;head involution;maintenance of polarity of embryonic epithelium |

| | | | | | | | | | | | |
|---|---|---|---|---|---|---|---|---|---|---|---|
| 2R | 9821791 | G | A | shot | 1.22E-04 | 0.443 | 0.449 | 0.497 | 0.368 | 9.30E-01 | microtubule cytoskeleton organization;astral microtubule;microtubule bundle formation;actin binding;calcium ion binding;protein binding;cytoskeleton;microtubule;adherens junction;microtubule-based process;negative regulation of microtubule depolymerization;cell cycle arrest;axonogenesis;sensory organ development;open tracheal system development;apposition of dorsal and ventral imaginal disc-derived wing surfaces;muscle organ development;microtubule binding;cytoskeletal protein binding;axon midline choice point recognition;muscle attachment;determination of muscle attachment site;mushroom body development;actin cytoskeleton organization;filopodium;growth cone;regulation of axon extension;oocyte fate determination;branch fusion |
| 2R | 9959741 | A | G | Prosap | 7.17E-05 | 0.616 | 0.621 | 0.598 | 0.694 | 2.63E-01 | protein binding;postsynaptic density;GKAP/Homer scaffold activity;postsynaptic density assembly |
| 2R | 1.1E+07 | T | C | CG30480 | 1.24E-04 | 0.270 | 0.316 | 0.248 | 0.391 | 9.02E-01 | |
| 2R | 1.1E+07 | T | A | mspo | 8.89E-05 | 0.216 | 0.191 | 0.223 | 0.093 | 1.09E-01 | regulation of myoblast fusion |
| 2R | 1.1E+07 | A | C | igl | 1.81E-05 | 0.517 | 0.485 | 0.504 | 0.440 | 5.71E-01 | calmodulin binding;cell cortex;myosin light chain binding |
| 2R | 1.1E+07 | A | T | trpm | 9.55E-05 | 0.489 | 0.436 | 0.367 | 0.473 | 5.34E-01 | zinc ion transmembrane transporter activity |
| 2R | 1.1E+07 | C | T | (CG8157) | 3.82E-05 | 0.192 | 0.202 | 0.160 | 0.225 | 1.48E-01 | NA |
| 2R | 1.2E+07 | C | T | shark | 8.40E-05 | 0.431 | 0.306 | 0.353 | 0.266 | 6.88E-01 | protein tyrosine kinase activity;non-membrane spanning protein tyrosine kinase activity;ATP binding;cytoplasm;cell cortex;protein phosphorylation;JNK cascade;eggshell chorion assembly;dorsal closure;dorsal closure |
| 2R | 1.2E+07 | T | C | (CG15711) | 6.43E-05 | 0.027 | 0.045 | 0.055 | 0.029 | 5.45E-01 | NA |

| | | | | | | | | | | | |
|---|---|---|---|---|---|---|---|---|---|---|---|
| 2R | 1.2E+07 | A | T | CG8311 | 4.49E-05 | 0.043 | 0.070 | 0.031 | 0.077 | 4.92E-01 | dolichol kinase activity;integral component of endoplasmic reticulum membrane;dolichyl monophosphate biosynthetic process |
| 2R | 1.3E+07 | A | T | CG8910 | 8.57E-05 | 0.097 | 0.099 | 0.066 | 0.131 | 1.52E-03 | zinc ion binding |
| 2R | 1.3E+07 | G | A | CG6796 | 2.58E-05 | 0.060 | 0.070 | 0.030 | 0.074 | 5.01E-01 | nucleic acid binding;asparagine-tRNA ligase activity;ATP binding;cytoplasm;asparaginyl-tRNA aminoacylation |
| 2R | 1.3E+07 | G | C | CG30460 | 5.00E-05 | 0.867 | 0.838 | 0.871 | 0.798 | NA | |
| 2R | 1.3E+07 | T | C | CG30460 | 1.36E-04 | 0.868 | 0.836 | 0.871 | 0.791 | NA | |
| 2R | 1.3E+07 | A | G | (CG15611) | 4.66E-05 | 0.031 | 0.063 | 0.080 | 0.026 | 5.00E-01 | NA |
| 2R | 1.3E+07 | C | T | mbl | 6.98E-05 | 0.400 | 0.353 | 0.233 | 0.372 | 2.17E-01 | regulation of alternative mRNA splicing |
| 2R | 1.4E+07 | C | A | elk | 6.46E-05 | 0.111 | 0.157 | 0.158 | 0.129 | 2.30E-01 | |
| 2R | 1.4E+07 | T | C | dpr13 | 6.36E-05 | 0.043 | 0.030 | 0.030 | 0.068 | NA | sensory perception of chemical stimulus |
| 2R | 1.4E+07 | G | A | (stau) | 5.70E-05 | 0.055 | 0.050 | 0.074 | 0.031 | 8.07E-01 | NA |
| 2R | 1.4E+07 | G | A | CG18537 | 1.79E-05 | 0.180 | 0.156 | 0.128 | 0.186 | 8.63E-01 | |
| 2R | 1.4E+07 | A | C | Pepck | 1.07E-04 | 0.385 | 0.436 | 0.341 | 0.498 | 3.57E-01 | phosphoenolpyruvate carboxykinase (GTP) activity;GTP binding;mitochondrion;gluconeogenesis |
| 2R | 1.5E+07 | C | A | Hs3st-A | 7.68E-06 | 0.305 | 0.322 | 0.227 | 0.365 | 9.54E-01 | [heparan sulfate]-glucosamine 3-sulfotransferase 1 activity |
| 2R | 1.5E+07 | A | C | (CG15115) | 7.57E-05 | 0.068 | 0.050 | 0.081 | 0.029 | 6.80E-01 | NA |
| 2R | 1.5E+07 | A | T | CG10081 | 6.12E-05 | 0.082 | 0.066 | 0.046 | 0.078 | 9.57E-02 | proteolysis;peptidase activity |
| 2R | 1.5E+07 | G | C | (sm) | 1.13E-04 | 0.116 | 0.089 | 0.088 | 0.052 | NA | NA |
| 2R | 1.7E+07 | G | C | (shg) | 1.37E-04 | 0.160 | 0.190 | 0.180 | 0.274 | 4.96E-02 | NA |
| 2R | 1.7E+07 | A | T | king-tubby | 1.35E-04 | 0.134 | 0.153 | 0.151 | 0.112 | 4.87E-01 | phosphatidylinositol binding;deactivation of rhodopsin mediated signaling;protein localization to cilium;glucose homeostasis |
| 2R | 1.7E+07 | C | A | Magi | 5.31E-05 | 0.281 | 0.405 | 0.390 | 0.311 | 2.54E-01 | guanylate kinase activity;membrane;Ral GTPase binding |
| 2R | 1.7E+07 | C | T | (Rgk3) | 1.45E-04 | 0.264 | 0.243 | 0.172 | 0.301 | NA | NA |
| 2R | 1.7E+07 | A | C | (Rgk3) | 1.26E-04 | 0.251 | 0.239 | 0.158 | 0.292 | NA | NA |
| 2R | 1.7E+07 | C | T | Rgk3 | 1.02E-04 | 0.306 | 0.359 | 0.273 | 0.381 | 2.28E-02 | GTP binding;obsolete GTP catabolic process;small GTPase mediated signal transduction;membrane |

| | | | | | | | | | | | |
|---|---|---|---|---|---|---|---|---|---|---|---|
| 2R | 1.7E+07 | G | C | CG10505 | 1.30E-04 | 0.147 | 0.084 | 0.062 | 0.126 | 2.11E-01 | transporter activity;ATP binding;response to zinc ion;ATPase activity |
| 2R | 1.7E+07 | T | G | Sdc | 1.34E-04 | 0.036 | 0.027 | 0.046 | 0.020 | 5.94E-01 | transmembrane signaling receptor activity;extracellular region;plasma membrane;focal adhesion;axon guidance;epithelial cell migration |
| 2R | 1.8E+07 | T | A | a | 8.94E-06 | 0.656 | 0.573 | 0.571 | 0.658 | NA | adherens junction;apical plasma membrane;compound eye development |
| 2R | 1.8E+07 | C | T | dve | 1.24E-04 | 0.288 | 0.249 | 0.305 | 0.178 | NA | negative regulation of transcription by RNA polymerase II;AT DNA binding;DNA-binding transcription factor activity;nucleus;regulation of transcription |
| 2R | 1.8E+07 | G | A | CG4554 | 1.10E-05 | 0.042 | 0.034 | 0.012 | 0.048 | 5.64E-01 | neurogenesis |
| 2R | 1.9E+07 | T | A | RpL23 | 3.26E-05 | 0.276 | 0.295 | 0.275 | 0.345 | 8.81E-01 | mitotic spindle elongation;structural constituent of ribosome;ribosome;translation;mitotic spindle organization;myosin binding;neurogenesis;cytosolic large ribosomal subunit |
| 2R | 2E+07 | C | T | CG9850 | 1.33E-04 | 0.084 | 0.052 | 0.095 | 0.029 | 1.71E-01 | metalloendopeptidase activity;proteolysis;cell population proliferation |
| 2R | 2E+07 | A | C | ytr | 8.56E-05 | 0.185 | 0.215 | 0.224 | 0.134 | 2.02E-01 | mRNA splicing |
| 2R | 2.1E+07 | T | C | emp | 2.18E-05 | 0.605 | 0.545 | 0.456 | 0.682 | 6.56E-01 | scavenger receptor activity;lysosome;plasma membrane;integral component of plasma membrane;defense response;cell adhesion;salivary gland cell autophagic cell death;autophagic cell death |
| 3L | 331062 | A | G | mthl9 | 8.76E-05 | 0.197 | 0.200 | 0.160 | 0.243 | 6.94E-01 | G protein-coupled receptor activity;response to stress;G protein-coupled receptor signaling pathway;determination of adult lifespan;integral component of membrane |
| 3L | 1269786 | C | G | CG9134 | 3.38E-05 | 0.068 | 0.055 | 0.073 | 0.026 | 4.91E-01 | carbohydrate binding |
| 3L | 1658473 | T | C | CG7971 | 5.94E-05 | 0.590 | 0.590 | 0.491 | 0.620 | 3.55E-01 | mRNA splicing |
| 3L | 2198846 | T | G | (CG8960) | 9.58E-05 | 0.873 | 0.866 | 0.854 | 0.900 | 5.45E-02 | NA |
| 3L | 2210365 | C | G | (CG15878) | 7.08E-05 | 0.634 | 0.607 | 0.738 | 0.538 | 2.26E-01 | NA |

| | | | | | | | | | | | |
|---|---|---|---|---|---|---|---|---|---|---|---|
| 3L | 2327513 | T | C | DmsR-1 | 7.35E-05 | 0.217 | 0.223 | 0.226 | 0.282 | NA | myosuppressin receptor activity;G protein-coupled receptor activity;neuropeptide receptor activity;adult locomotory behavior |
| 3L | 2964616 | C | A | (CG34025) | 1.67E-05 | 0.115 | 0.131 | 0.083 | 0.146 | NA | NA |
| 3L | 3476925 | C | A | (CG32267) | 2.14E-05 | 0.035 | 0.040 | 0.025 | 0.058 | 9.71E-01 | NA |
| 3L | 3844675 | A | G | (Awh) | 7.39E-05 | 0.832 | 0.777 | 0.830 | 0.760 | 6.00E-01 | NA |
| 3L | 4404707 | A | G | Syx17 | 5.37E-05 | 0.589 | 0.581 | 0.667 | 0.500 | 1.52E-02 | SNARE binding;autophagosome membrane;SNAP receptor activity;plasma membrane;endoplasmic reticulum to Golgi vesicle-mediated transport;neurotransmitter secretion;synaptic vesicle docking;vesicle-mediated transport;SNARE complex;neuron cellular homeostasis;endoplasmic reticulum-Golgi intermediate compartment organization;autophagosome maturation |
| 3L | 4842109 | C | A | (CG13707) | 2.50E-05 | 0.923 | 0.926 | 0.924 | 0.953 | 7.18E-01 | NA |
| 3L | 5389566 | G | C | (Ir64a) | 9.40E-05 | 0.600 | 0.592 | 0.483 | 0.606 | NA | NA |
| 3L | 6590431 | A | G | ndl | 1.23E-04 | 0.117 | 0.099 | 0.153 | 0.105 | NA | serine-type endopeptidase activity;extracellular region;proteolysis;eggshell chorion assembly;oocyte dorsal/ventral axis specification;maternal specification of dorsal/ventral axis |
| 3L | 7335246 | G | A | CG8596 | 3.16E-05 | 0.109 | 0.077 | 0.099 | 0.141 | 8.79E-05 | integral component of membrane;transmembrane transport |
| 3L | 7454495 | T | C | (CG42660) | 1.33E-04 | 0.081 | 0.039 | 0.048 | 0.020 | 3.66E-01 | NA |
| 3L | 7639899 | A | T | (CG32373) | 4.57E-05 | 0.331 | 0.283 | 0.212 | 0.358 | 1.14E-01 | NA |
| 3L | 7951378 | T | C | (exex) | 1.02E-04 | 0.051 | 0.034 | 0.051 | 0.022 | 6.02E-01 | NA |
| 3L | 8284093 | G | A | CG7201 | 8.33E-05 | 0.074 | 0.071 | 0.088 | 0.033 | 4.19E-01 | |
| 3L | 8318762 | G | C | (CG34461) | 4.52E-05 | 0.462 | 0.342 | 0.432 | 0.317 | 8.96E-01 | NA |
| 3L | 8606593 | T | G | CG6282 | 7.18E-05 | 0.224 | 0.237 | 0.115 | 0.200 | 9.25E-01 | cytoplasm;lipid metabolic process;integral component of membrane;oxidoreductase activity |
| 3L | 8688265 | G | A | (h) | 1.04E-04 | 0.046 | 0.051 | 0.052 | 0.022 | 3.73E-01 | NA |

| | | | | | | | | | | | |
|---|---|---|---|---|---|---|---|---|---|---|---|
| 3L | 8843663 | C | T | dally | 1.05E-04 | 0.210 | 0.224 | 0.304 | 0.187 | NA | protein binding;extracellular region;NA;transforming growth factor beta receptor signaling pathway;nervous system development;axonogenesis;wing disc morphogenesis;imaginal disc-derived wing morphogenesis;imaginal disc-derived leg morphogenesis;motor neuron axon guidance;sensory organ boundary specification;decapentaplegic signaling pathway;G2/MI transition of meiotic cell cycle;chaeta morphogenesis;imaginal disc-derived wing vein morphogenesis;cell surface;membrane;Wnt signaling pathway;extrinsic component of plasma membrane;chaeta development;regulation of BMP signaling pathway;positive regulation of BMP signaling pathway;germ-line stem cell population maintenance;wing disc development;regulation of multicellular organism growth;heparan sulfate proteoglycan binding;regulation of imaginal disc growth;female germ-line stem cell asymmetric division;wing disc dorsal/ventral pattern formation;compound eye development;dendrite morphogenesis;regulation of cell cycle;negative regulation of semaphorin-plexin signaling pathway |
| 3L | 9368910 | C | G | CG4461 | 5.90E-05 | 0.789 | 0.797 | 0.732 | 0.803 | 6.83E-01 | response to heat |
| 3L | 9575690 | C | T | CG42673 | 1.33E-05 | 0.403 | 0.453 | 0.504 | 0.426 | 1.93E-01 | |
| 3L | 9593255 | T | A | CG3222 | 1.36E-04 | 0.579 | 0.630 | 0.608 | 0.548 | 5.60E-01 | |
| 3L | 9769486 | G | T | CG8177 | 1.49E-04 | 0.048 | 0.062 | 0.030 | 0.080 | 8.97E-01 | inorganic anion exchanger activity;anion transport;anion:anion antiporter activity;integral component of membrane |
| 3L | 1E+07 | T | C | (dpr10) | 5.84E-06 | 0.058 | 0.038 | 0.027 | 0.050 | 3.32E-03 | NA |
| 3L | 1E+07 | T | G | (dpr10) | 3.31E-05 | 0.056 | 0.041 | 0.027 | 0.052 | 2.28E-03 | NA |

| | | | | | | | | | | | |
|---|---|---|---|---|---|---|---|---|---|---|---|
| 3L | 1E+07 | C | T | A2bp1 | 1.33E-05 | 0.055 | 0.062 | 0.079 | 0.027 | 1.60E-01 | nucleotide binding;mRNA binding;nucleus;cytoplasm;nervous system development;imaginal disc-derived wing vein specification;transcription factor binding;transcription regulatory region DNA binding;positive regulation of transcription |
| 3L | 1.1E+07 | G | A | (NijA) | 1.28E-04 | 0.344 | 0.365 | 0.287 | 0.423 | 5.81E-02 | NA |
| 3L | 1.1E+07 | T | A | (Aps) | 1.01E-04 | 0.404 | 0.472 | 0.459 | 0.532 | 6.64E-01 | NA |
| 3L | 1.1E+07 | G | A | (CG34050) | 5.58E-05 | 0.419 | 0.455 | 0.351 | 0.485 | NA | NA |
| 3L | 1.1E+07 | T | G | CG7638 | 9.40E-05 | 0.086 | 0.088 | 0.151 | 0.096 | 9.05E-02 | |
| 3L | 1.1E+07 | G | A | GlcAT-P | 1.09E-04 | 0.032 | 0.031 | 0.011 | 0.051 | 6.41E-01 | glycosphingolipid biosynthetic process;peripheral nervous system development;glycoprotein biosynthetic process;galactosylgalactosylxylosylprotein 3-beta-glucuronosyltransferase activity;glucuronosyltransferase activity;membrane;proteoglycan biosynthetic process;N-acetyllactosamine beta-1 |
| 3L | 1.1E+07 | G | A | (CG6168) | 1.85E-05 | 0.115 | 0.131 | 0.256 | 0.107 | 7.66E-01 | NA |
| 3L | 1.1E+07 | C | A | (CG6163) | 1.41E-04 | 0.171 | 0.185 | 0.147 | 0.195 | 4.91E-01 | NA |
| 3L | 1.2E+07 | T | C | Mob2 | 9.70E-05 | 0.068 | 0.100 | 0.050 | 0.134 | 9.23E-01 | cell morphogenesis;protein binding;cytoplasm;plasma membrane;learning or memory;long-term memory;olfactory learning;rhabdomere;protein kinase binding;rhabdomere development;negative regulation of synaptic growth at neuromuscular junction;cell hair;cell periphery |

| | | | | | | | | | | | |
|---|---|---|---|---|---|---|---|---|---|---|---|
| 3L | 1.2E+07 | T | G | Nrx-IV | 1.34E-04 | 0.858 | 0.867 | 0.877 | 0.768 | 3.76E-05 | heart process;transmembrane signaling receptor activity;plasma membrane;integral component of plasma membrane;septate junction;pleated septate junction;establishment or maintenance of cell polarity;dorsal closure;synaptic target recognition;establishment of blood-nerve barrier;protein localization;axon ensheathment;integral component of membrane;synaptic vesicle targeting;synaptic vesicle docking;septate junction assembly;nerve maturation;regulation of tube size |
| 3L | 1.3E+07 | A | C | sowah | 5.27E-05 | 0.461 | 0.459 | 0.529 | 0.419 | 9.54E-01 | |
| 3L | 1.3E+07 | G | C | CG32111 | 5.53E-05 | 0.076 | 0.032 | 0.052 | 0.026 | 6.34E-01 | |
| 3L | 1.3E+07 | C | G | (mirr) | 8.82E-06 | 0.034 | 0.037 | 0.026 | 0.052 | 1.08E-01 | NA |
| 3L | 1.3E+07 | C | T | (mirr) | 1.07E-04 | 0.177 | 0.168 | 0.108 | 0.189 | 2.98E-02 | NA |
| 3L | 1.3E+07 | G | A | (CG10943) | 7.74E-05 | 0.027 | 0.025 | 0.044 | 0.096 | 6.44E-01 | NA |
| 3L | 1.3E+07 | C | T | (CG11262) | 3.30E-05 | 0.557 | 0.539 | 0.443 | 0.588 | NA | NA |
| 3L | 1.3E+07 | T | A | CG17672 | 6.16E-05 | 0.066 | 0.058 | 0.010 | 0.064 | 6.54E-01 | |
| 3L | 1.3E+07 | G | A | (CR43913) | 2.79E-05 | 0.215 | 0.249 | 0.142 | 0.272 | 4.10E-01 | NA |
| 3L | 1.3E+07 | G | T | (CR43912) | 8.87E-05 | 0.042 | 0.031 | 0.072 | 0.026 | 3.17E-01 | NA |
| 3L | 1.4E+07 | A | T | Sox21b | 1.39E-04 | 0.101 | 0.154 | 0.201 | 0.108 | 8.62E-01 | DNA-binding transcription factor activity;nucleus;regulation of transcription |
| 3L | 1.5E+07 | C | A | bbg | 1.27E-04 | 0.063 | 0.050 | 0.021 | 0.046 | 9.38E-01 | nucleus;plasma membrane;border follicle cell migration;imaginal disc development |
| 3L | 1.5E+07 | C | A | (BobA) | 1.35E-04 | 0.027 | 0.035 | 0.013 | 0.054 | 8.84E-01 | NA |
| 3L | 1.5E+07 | A | T | (Tollo) | 1.04E-04 | 0.148 | 0.189 | 0.254 | 0.155 | 9.53E-01 | NA |
| 3L | 1.5E+07 | A | T | (Tollo) | 4.77E-05 | 0.141 | 0.188 | 0.222 | 0.126 | 6.97E-01 | NA |
| 3L | 1.5E+07 | G | T | (CR43625) | 1.33E-04 | 0.581 | 0.548 | 0.418 | 0.584 | 8.15E-01 | NA |
| 3L | 1.6E+07 | T | C | (CG34451) | 1.12E-04 | 0.757 | 0.754 | 0.771 | 0.748 | 4.20E-01 | NA |
| 3L | 1.6E+07 | C | A | sff | 8.40E-05 | 0.126 | 0.136 | 0.155 | 0.084 | 8.88E-01 | protein serine/threonine kinase activity;ATP binding;protein phosphorylation;protein N-linked glycosylation;neuromuscular junction development |
| 3L | 1.6E+07 | G | T | CG13070 | 9.09E-05 | 0.348 | 0.316 | 0.340 | 0.369 | NA | |

| | | | | | | | | | | | |
|---|---|---|---|---|---|---|---|---|---|---|---|
| 3L | 1.6E+07 | C | T | CG33158 | 3.73E-05 | 0.075 | 0.059 | 0.074 | 0.028 | 7.91E-02 | translation elongation factor activity;GTPase activity;GTP binding;eukaryotic translation initiation factor 2 complex;translational elongation |
| 3L | 1.6E+07 | T | C | CG33158 | 1.21E-05 | 0.077 | 0.061 | 0.077 | 0.029 | 7.77E-02 | translation elongation factor activity;GTPase activity;GTP binding;eukaryotic translation initiation factor 2 complex;translational elongation |
| 3L | 1.8E+07 | G | A | (CG5290) | 2.51E-05 | 0.194 | 0.174 | 0.209 | 0.130 | 9.54E-04 | NA |
| 3L | 1.8E+07 | T | A | (grim) | 4.02E-05 | 0.958 | 0.951 | 0.971 | 0.939 | 2.89E-01 | NA |
| 3L | 1.9E+07 | C | A | (CG32204) | 1.09E-04 | 0.053 | 0.108 | 0.125 | 0.074 | NA | NA |
| 3L | 1.9E+07 | T | C | (Spn75F) | 1.33E-04 | 0.039 | 0.047 | 0.071 | 0.023 | 6.55E-02 | NA |
| 3L | 1.9E+07 | C | T | CG32206 | 1.26E-04 | 0.077 | 0.082 | 0.056 | 0.029 | 3.16E-01 | Wnt-protein binding;Wnt-activated receptor activity;Wnt signaling pathway involved in dorsal/ventral axis specification;lateral inhibition;canonical Wnt signaling pathway |
| 3L | 2E+07 | A | G | CG42674 | 1.05E-04 | 0.229 | 0.196 | 0.179 | 0.228 | 4.95E-04 | Rho guanyl-nucleotide exchange factor activity;cell adhesion;imaginal disc-derived leg morphogenesis;regulation of cell shape;positive regulation of Rho protein signal transduction |
| 3L | 2.1E+07 | A | G | (CR43929) | 1.52E-04 | 0.136 | 0.103 | 0.177 | 0.096 | 7.73E-01 | NA |
| 3L | 2.2E+07 | G | T | (mir-4942) | 5.67E-05 | 0.063 | 0.217 | 0.173 | 0.273 | 2.19E-01 | NA |

| | | | | | | | | | | | |
|---|---|---|---|---|---|---|---|---|---|---|---|
| 3L | 2.3E+07 | G | A | alpha-Cat | 1.33E-04 | 0.026 | 0.025 | 0.091 | 0.052 | 1.58E-01 | actin binding;structural molecule activity;protein binding;plasma membrane;adherens junction;cell-cell adherens junction;spot adherens junction;zonula adherens;cytoskeletal anchoring at plasma membrane;cell adhesion;establishment or maintenance of cell polarity;cytoskeletal protein binding;actin cytoskeleton;catenin complex;cell-substrate junction;oocyte localization involved in germarium-derived egg chamber formation;adherens junction organization;cadherin binding;head morphogenesis |
| 3R | 1262668 | C | A | CG14669 | 4.50E-05 | 0.281 | 0.233 | 0.278 | 0.214 | 1.15E-04 | GTP binding;obsolete GTP catabolic process;small GTPase mediated signal transduction;membrane |
| 3R | 1892799 | G | T | (CG15580) | 8.55E-05 | 0.407 | 0.353 | 0.315 | 0.373 | 4.86E-01 | NA |
| 3R | 2456272 | T | C | sunz | 1.51E-04 | 0.275 | 0.290 | 0.215 | 0.301 | 3.58E-01 | calcium ion binding;male meiotic nuclear division |
| 3R | 5950686 | A | G | knk | 1.22E-04 | 0.053 | 0.116 | 0.156 | 0.079 | 2.48E-04 | embryonic epithelial tube formation;chitin biosynthetic process;cuticle chitin biosynthetic process;terminal region determination;open tracheal system development;torso signaling pathway;chitin-based embryonic cuticle biosynthetic process;regulation of tube size |
| 3R | 6250938 | C | A | (CG6345) | 8.76E-06 | 0.582 | 0.562 | 0.518 | 0.608 | 2.02E-01 | NA |
| 3R | 7105520 | G | C | CG31386 | 1.37E-04 | 0.580 | 0.410 | 0.414 | 0.532 | 2.68E-01 | |
| 3R | 7634750 | G | A | ClC-a | 2.83E-05 | 0.026 | 0.039 | 0.011 | 0.045 | 4.50E-01 | voltage-gated chloride channel activity;chloride channel activity;chloride transport;actin cytoskeleton;membrane;adenyl nucleotide binding;transmembrane transport |
| 3R | 7982089 | G | C | dpr15 | 8.24E-05 | 0.097 | 0.175 | 0.192 | 0.140 | 1.28E-02 | sensory perception of chemical stimulus |

| | | | | | | | | | | | |
|---|---|---|---|---|---|---|---|---|---|---|---|
| 3R | 7990369 | G | C | Cpn | 5.55E-05 | 0.041 | 0.056 | 0.055 | 0.013 | 1.65E-01 | calcium ion binding;nucleus;cytoplasm;mitochondrion;rough endoplasmic reticulum;rhabdomere;axon;rhabdomere development;cell body;sequestering of calcium ion |
| 3R | 8426065 | G | A | (Octbeta2R) | 5.94E-05 | 0.205 | 0.175 | 0.143 | 0.090 | 1.07E-02 | NA |
| 3R | 9224993 | T | C | CG9796 | 2.64E-05 | 0.138 | 0.109 | 0.046 | 0.078 | 7.87E-02 | |
| 3R | 9820259 | C | G | rdx | 8.81E-05 | 0.261 | 0.267 | 0.263 | 0.189 | 6.85E-01 | protein binding;nucleus;cellularization;protein ubiquitination;regulation of proteolysis;protein destabilization;establishment of ommatidial planar polarity;negative regulation of protein import into nucleus;protein homodimerization activity;positive regulation of apoptotic process;negative regulation of smoothened signaling pathway;positive regulation of JNK cascade;lateral inhibition;eye morphogenesis |
| 3R | 1E+07 | T | C | (cv-c) | 1.42E-04 | 0.227 | 0.240 | 0.200 | 0.293 | 9.87E-01 | NA |
| 3R | 1.1E+07 | A | T | Neu3 | 1.21E-04 | 0.035 | 0.023 | 0.011 | 0.033 | 2.02E-01 | |
| 3R | 1.1E+07 | T | C | (btsz) | 1.37E-04 | 0.066 | 0.086 | 0.075 | 0.151 | 4.19E-01 | NA |
| 3R | 1.1E+07 | C | A | (CG3837) | 3.19E-05 | 0.096 | 0.114 | 0.100 | 0.046 | 2.26E-01 | NA |
| 3R | 1.1E+07 | T | G | (CG43335) | 5.07E-05 | 0.433 | 0.596 | 0.672 | 0.554 | 2.38E-02 | NA |
| 3R | 1.1E+07 | G | C | (pxb) | 1.88E-05 | 0.017 | 0.026 | 0.041 | 0.010 | 6.69E-01 | NA |
| 3R | 1.2E+07 | C | T | alpha-Man-IIb | 1.16E-04 | 0.078 | 0.093 | 0.132 | 0.089 | 5.63E-01 | Golgi membrane;alpha-mannosidase activity;mannosyl-oligosaccharide 1 |
| 3R | 1.2E+07 | T | G | ss | 7.25E-05 | 0.081 | 0.084 | 0.054 | 0.107 | 8.52E-05 | negative regulation of transcription by RNA polymerase II;DNA binding;DNA-binding transcription factor activity;NA;obsolete signal transducer activity;nucleus;regulation of transcription |
| 3R | 1.3E+07 | A | T | msa | 5.21E-05 | 0.317 | 0.335 | 0.295 | 0.390 | 2.89E-02 | |
| 3R | 1.3E+07 | T | A | (Mur89F) | 1.47E-04 | 0.030 | 0.066 | 0.097 | 0.027 | NA | NA |
| 3R | 1.3E+07 | C | A | (CG31262) | 1.33E-04 | 0.298 | 0.230 | 0.175 | 0.293 | 4.45E-01 | NA |
| 3R | 1.4E+07 | C | T | CG18012 | 5.36E-06 | 0.270 | 0.296 | 0.311 | 0.225 | 4.68E-03 | protein glycosylation;beta-1 |

| | | | | | | | | | | | |
|---|---|---|---|---|---|---|---|---|---|---|---|
| 3R | 1.4E+07 | T | C | (htl) | 1.01E-05 | 0.876 | 0.884 | 0.947 | 0.855 | 4.11E-02 | NA |
| 3R | 1.4E+07 | G | A | CG14316 | 3.72E-05 | 0.191 | 0.159 | 0.137 | 0.079 | 7.39E-01 | biological_process |
| 3R | 1.4E+07 | G | T | CG7156 | 1.25E-04 | 0.085 | 0.150 | 0.130 | 0.189 | 1.57E-04 | protein kinase activity;ATP binding;protein phosphorylation;phosphatidylinositol binding;neuron projection morphogenesis |
| 3R | 1.5E+07 | T | A | CG42613 | 1.55E-05 | 0.193 | 0.248 | 0.300 | 0.245 | NA | biological_process |
| 3R | 1.5E+07 | A | G | CG5316 | 1.50E-04 | 0.169 | 0.137 | 0.210 | 0.148 | 8.88E-01 | single strand break repair;mRNA splicing |
| 3R | 1.6E+07 | C | T | mun | 9.40E-05 | 0.389 | 0.394 | 0.300 | 0.412 | 1.39E-01 | |
| 3R | 1.7E+07 | T | G | lbl | 3.71E-05 | 0.735 | 0.655 | 0.658 | 0.771 | 2.22E-01 | transcription regulatory region sequence-specific DNA binding;DNA-binding transcription factor activity;nucleus;regulation of transcription |
| 3R | 1.9E+07 | A | G | CG13830 | 1.22E-04 | 0.339 | 0.311 | 0.354 | 0.255 | NA | calcium ion binding;NA;signal transduction |
| 3R | 1.9E+07 | T | C | CG6763 | 8.94E-05 | 0.552 | 0.532 | 0.493 | 0.684 | 1.49E-02 | metalloendopeptidase activity;proteolysis;zinc ion binding;meprin A complex |
| 3R | 2E+07 | A | G | mbc | 2.21E-05 | 0.795 | 0.713 | 0.741 | 0.645 | NA | cell morphogenesis;ruffle;protein binding;cytoplasm;cytoskeleton organization;border follicle cell migration;dorsal closure;central nervous system development;skeletal muscle tissue development;myoblast fusion;larval visceral muscle development;muscle attachment;extrinsic component of plasma membrane;actin cytoskeleton organization;Rac guanyl-nucleotide exchange factor activity;NA;cell competition in a multicellular organism;anterior Malpighian tubule development |
| 3R | 2E+07 | T | A | (Esyt2) | 1.03E-04 | 0.323 | 0.320 | 0.315 | 0.219 | 7.54E-01 | NA |
| 3R | 2E+07 | G | A | CG13624 | 1.00E-04 | 0.257 | 0.313 | 0.196 | 0.333 | 3.76E-02 | DNA-binding transcription factor activity;regulation of transcription |
| 3R | 2.1E+07 | T | A | CG31370 | 4.07E-05 | 0.090 | 0.065 | 0.109 | 0.052 | 7.04E-01 | transferase activity |
| 3R | 2.2E+07 | T | A | CG4743 | 1.96E-05 | 0.024 | 0.038 | 0.050 | 0.013 | 2.04E-01 | mitochondrial envelope;transmembrane transporter activity;transmembrane transport |
| 3R | 2.2E+07 | T | C | malpha | 8.38E-05 | 0.410 | 0.364 | 0.475 | 0.350 | 1.61E-03 | |

| | | | | | | | | | | | |
|---|---|---|---|---|---|---|---|---|---|---|---|
| 3R | 2.2E+07 | C | T | (CG6073) | 1.15E-04 | 0.371 | 0.418 | 0.368 | 0.473 | 1.04E-01 | NA |
| 3R | 2.2E+07 | T | A | (CG6073) | 7.44E-05 | 0.371 | 0.415 | 0.368 | 0.476 | 1.55E-01 | NA |
| 3R | 2.3E+07 | G | C | CG14253 | 7.63E-05 | 0.524 | 0.608 | 0.516 | 0.666 | 2.81E-01 | |
| 3R | 2.3E+07 | C | T | CG5611 | 1.09E-04 | 0.142 | 0.186 | 0.070 | 0.200 | 3.39E-01 | metabolic process |
| 3R | 2.4E+07 | A | T | CG34353 | 1.51E-04 | 0.086 | 0.095 | 0.180 | 0.091 | 1.99E-01 | gravitaxis |
| 3R | 2.5E+07 | G | A | CG10000 | 5.00E-05 | 0.074 | 0.091 | 0.127 | 0.065 | 5.20E-03 | polypeptide N-acetylgalactosaminyltransferase activity;Golgi stack;oligosaccharide biosynthetic process;multicellular organism reproduction |
| 3R | 2.5E+07 | C | A | CG10000 | 6.19E-05 | 0.037 | 0.020 | 0.056 | 0.010 | 1.01E-01 | polypeptide N-acetylgalactosaminyltransferase activity;Golgi stack;oligosaccharide biosynthetic process;multicellular organism reproduction |
| 3R | 2.5E+07 | G | C | (CG12558) | 1.94E-05 | 0.526 | 0.408 | 0.401 | 0.522 | 2.20E-03 | NA |
| 3R | 2.5E+07 | G | A | CG14521 | 7.81E-05 | 0.052 | 0.033 | 0.012 | 0.049 | 9.35E-01 | |
| 3R | 2.5E+07 | A | C | CG11897 | 1.48E-04 | 0.469 | 0.474 | 0.421 | 0.526 | 3.50E-01 | ATP binding;xenobiotic transmembrane transporting ATPase activity;drug transmembrane transporter activity;integral component of membrane;transmembrane transport |
| 3R | 2.5E+07 | T | A | (Cnx99A) | 8.52E-05 | 0.082 | 0.083 | 0.037 | 0.110 | 3.16E-01 | NA |
| 3R | 2.5E+07 | G | C | DopR2 | 1.43E-04 | 0.486 | 0.539 | 0.491 | 0.625 | 6.78E-01 | |
| 3R | 2.6E+07 | G | A | ncd | 1.49E-04 | 0.148 | 0.145 | 0.093 | 0.208 | 3.16E-02 | mitotic spindle elongation;meiotic spindle organization;microtubule bundle formation;microtubule motor activity;ATP binding;spindle;kinesin complex;minus-end kinesin complex;microtubule-based movement;spindle organization;mitotic spindle organization;spindle assembly involved in female meiosis;chromosome segregation;mitotic centrosome separation;microtubule binding;ATP-dependent microtubule motor activity |
| 3R | 2.6E+07 | T | C | CG1983 | 1.19E-04 | 0.202 | 0.217 | 0.169 | 0.222 | 1.25E-01 | pyridoxal phosphate binding |
| 3R | 2.6E+07 | G | T | hdc | 1.40E-04 | 0.236 | 0.243 | 0.340 | 0.206 | NA | cytoplasm;open tracheal system development;terminal branching |

| | | | | | | | | | | | |
|---|---|---|---|---|---|---|---|---|---|---|---|
| 3R | 2.6E+07 | T | A | PH4alphaEFB | 2.92E-05 | 0.051 | 0.040 | 0.082 | 0.025 | NA | procollagen-proline 4-dioxygenase activity;iron ion binding;endoplasmic reticulum;procollagen-proline 4-dioxygenase complex;oxidoreductase activity |
| 3R | 2.6E+07 | C | T | (CG31013) | 2.13E-05 | 0.050 | 0.094 | 0.073 | 0.151 | 1.69E-02 | NA |
| 3R | 2.7E+07 | C | T | (CG15545) | 2.89E-05 | 0.238 | 0.261 | 0.168 | 0.288 | 1.27E-01 | NA |
| 3R | 2.7E+07 | G | A | (CG12071) | 2.31E-05 | 0.447 | 0.444 | 0.387 | 0.520 | 6.72E-01 | NA |
| 3R | 2.7E+07 | C | T | (CG15550) | 1.51E-04 | 0.075 | 0.088 | 0.050 | 0.081 | 3.48E-01 | NA |
| 3R | 2.7E+07 | G | A | CG12054 | 4.54E-05 | 0.061 | 0.046 | 0.028 | 0.060 | 3.03E-02 | nucleic acid binding;metal ion binding |
| 3R | 2.7E+07 | C | T | (CG1607) | 2.29E-05 | 0.536 | 0.520 | 0.399 | 0.548 | 5.17E-04 | NA |
| 3R | 2.7E+07 | A | C | Gprk2 | 5.11E-05 | 0.753 | 0.786 | 0.799 | 0.720 | 3.86E-01 | regulation of antimicrobial peptide biosynthetic process;apical constriction involved in gastrulation;protein serine/threonine kinase activity;G protein-coupled receptor kinase activity;ATP binding;cytoplasm;plasma membrane;protein phosphorylation;signal transduction;G protein-coupled receptor signaling pathway;smoothened signaling pathway;vitellogenesis;imaginal disc-derived wing vein specification;regulation of G protein-coupled receptor signaling pathway;regulation of smoothened signaling pathway;regulation of Toll signaling pathway;embryo development;termination of G protein-coupled receptor signaling pathway;positive regulation of cAMP-mediated signaling;negative regulation of smoothened signaling pathway;defense response to Gram-positive bacterium |
| 3R | 2.7E+07 | G | A | RpL6 | 4.20E-06 | 0.424 | 0.387 | 0.375 | 0.430 | 1.76E-02 | mitotic spindle elongation;structural constituent of ribosome;ribosome;translation;mitotic spindle organization;cytosolic large ribosomal subunit;NA;centrosome duplication |
| X | 343943 | C | A | CG32816 | 1.31E-04 | 0.300 | 0.225 | 0.195 | 0.318 | 9.93E-02 | |

| | | | | | | | | | | | |
|---|---|---|---|---|---|---|---|---|---|---|---|
| X | 1420447 | T | C | Mur2B | 5.20E-05 | 0.041 | 0.049 | 0.051 | 0.013 | 4.27E-01 | extracellular matrix structural constituent;extracellular region;chitin metabolic process;chitin binding;extracellular matrix;chorion |
| X | 2381099 | A | G | trol | 4.18E-05 | 0.312 | 0.289 | 0.205 | 0.358 | 2.51E-01 | NA;basement membrane;lipid droplet;defasciculation of motor neuron axon;motor neuron axon guidance;maintenance of epithelial cell apical/basal polarity;asymmetric neuroblast division;response to anesthetic;positive regulation of semaphorin-plexin signaling pathway |
| X | 2461125 | A | G | Pdfr | 3.54E-05 | 0.022 | 0.045 | 0.012 | 0.056 | 8.26E-01 | G protein-coupled receptor activity;calcitonin receptor activity;integral component of plasma membrane;G protein-coupled receptor signaling pathway;neuropeptide signaling pathway;circadian rhythm;neuropeptide receptor activity;NA;integral component of membrane;gravitaxis;circadian sleep/wake cycle;regulation of circadian sleep/wake cycle;neuron projection;neuronal cell body;locomotor rhythm;circadian behavior |
| X | 2602978 | C | A | CG2681 | 8.94E-05 | 0.313 | 0.335 | 0.376 | 0.317 | 7.48E-02 | ubiquitin-protein transferase activity;nucleus;ubiquitin-dependent protein catabolic process;multicellular organism development;zinc ion binding;protein ubiquitination |
| X | 3326693 | G | A | CG12535 | 2.43E-06 | 0.026 | 0.018 | 0.013 | 0.045 | 2.64E-01 | |
| X | 5495935 | G | T | Vsx1 | 1.43E-04 | 0.055 | 0.038 | 0.013 | 0.053 | 2.81E-01 | negative regulation of transcription by RNA polymerase II;optic lobe placode development;DNA-binding transcription factor activity;nucleus;phagocytosis;sequence-specific DNA binding;positive regulation of neural precursor cell proliferation |
| X | 5891394 | C | T | (CG5966) | 9.11E-05 | 0.075 | 0.088 | 0.031 | 0.053 | 7.50E-01 | NA |

| | | | | | | | | | | | |
|---|---|---|---|---|---|---|---|---|---|---|---|
| X | 5929941 | A | G | rux | 9.56E-05 | 0.332 | 0.381 | 0.393 | 0.305 | 2.71E-01 | mitotic cell cycle;nucleus;regulation of mitotic nuclear division;regulation of exit from mitosis;eye-antennal disc morphogenesis;regulation of meiotic nuclear division;compound eye development;regulation of cell cycle |
| X | 6549545 | G | A | l(1)G0148 | 1.02E-04 | 0.965 | 0.983 | 0.988 | 0.956 | 7.28E-02 | protein serine/threonine kinase activity;obsolete signal transducer |
| X | 6689046 | T | C | C3G | 6.98E-05 | 0.035 | 0.057 | 0.105 | 0.048 | NA | Ras guanyl-nucleotide exchange factor activity;intracellular;Ras protein signal transduction;somatic muscle development;muscle attachment;Rap guanyl-nucleotide exchange factor activity;NA;sarcomere organization |
| X | 8907108 | A | G | rdgA | 1.36E-04 | 0.077 | 0.052 | 0.092 | 0.023 | 4.05E-01 | NAD+ kinase activity;diacylglycerol kinase activity;microtubule associated complex;phosphatidic acid biosynthetic process;phosphatidylinositol biosynthetic process;actin filament organization;protein kinase C-activating G protein-coupled receptor signaling pathway;visual perception;phototransduction;sensory perception of sound;sensory perception of smell;metabolic process;membrane;rhodopsin mediated signaling pathway;deactivation of rhodopsin mediated signaling;phosphorylation;diacylglycerol binding;intracellular signal transduction;thermotaxis;photoreceptor cell maintenance;lipid phosphorylation |

| | | | | | | | | | | | |
|---|---|---|---|---|---|---|---|---|---|---|---|
| X | 8907110 | T | C | rdgA | 8.00E-05 | 0.062 | 0.053 | 0.092 | 0.023 | 3.89E-01 | NAD+ kinase activity;diacylglycerol kinase activity;microtubule associated complex;phosphatidic acid biosynthetic process;phosphatidylinositol biosynthetic process;actin filament organization;protein kinase C-activating G protein-coupled receptor signaling pathway;visual perception;phototransduction;sensory perception of sound;sensory perception of smell;metabolic process;membrane;rhodopsin mediated signaling pathway;deactivation of rhodopsin mediated signaling;phosphorylation;diacylglycerol binding;intracellular signal transduction;thermotaxis;photoreceptor cell maintenance;lipid phosphorylation |
| X | 8907115 | G | A | rdgA | 1.40E-04 | 0.083 | 0.052 | 0.093 | 0.023 | 5.38E-01 | NAD+ kinase activity;diacylglycerol kinase activity;microtubule associated complex;phosphatidic acid biosynthetic process;phosphatidylinositol biosynthetic process;actin filament organization;protein kinase C-activating G protein-coupled receptor signaling pathway;visual perception;phototransduction;sensory perception of sound;sensory perception of smell;metabolic process;membrane;rhodopsin mediated signaling pathway;deactivation of rhodopsin mediated signaling;phosphorylation;diacylglycerol binding;intracellular signal transduction;thermotaxis;photoreceptor cell maintenance;lipid phosphorylation |
| X | 9803147 | C | G | (CG1986) | 1.15E-04 | 0.143 | 0.128 | 0.057 | 0.105 | 3.70E-01 | NA |

| | | | | | | | | | | | |
|---|---|---|---|---|---|---|---|---|---|---|---|
| X | 1E+07 | C | T | CG1628 | 1.46E-04 | 0.035 | 0.039 | 0.078 | 0.042 | 6.37E-01 | L-ornithine transmembrane transporter activity;NA;amino acid transmembrane transport;mitochondrial envelope;mitochondrial inner membrane;amino acid transmembrane transporter activity;transmembrane transporter activity |
| X | 1.2E+07 | A | T | CR32661 | 7.83E-05 | 0.099 | 0.077 | 0.027 | 0.095 | 4.13E-01 | |
| X | 1.4E+07 | T | A | RNA:S474:12E | 1.16E-04 | 0.115 | 0.114 | 0.142 | 0.087 | 4.31E-01 | NA |
| X | 1.5E+07 | G | A | (Grip128) | 7.12E-05 | 0.290 | 0.273 | 0.222 | 0.306 | 2.48E-01 | NA |
| X | 1.7E+07 | C | A | (CG12998) | 9.04E-05 | 0.019 | 0.014 | 0.014 | 0.052 | 4.44E-01 | NA |
| X | 1.7E+07 | A | G | 3-H2-CR43491 | 3.95E-05 | 0.513 | 0.530 | 0.589 | 0.500 | 8.13E-01 | NA |
| X | 1.8E+07 | A | C | (RhoGAPp190) | 1.20E-05 | 0.251 | 0.263 | 0.199 | 0.317 | 7.94E-01 | NA |
| X | 1.8E+07 | A | G | CG15056 | 1.34E-04 | 0.026 | 0.021 | 0.012 | 0.043 | NA | G2/M transition of mitotic cell cycle;ubiquitin-protein transferase activity;ubiquitin-dependent protein catabolic process;SCF ubiquitin ligase complex;positive regulation of BMP signaling pathway;SCF-dependent proteasomal ubiquitin-dependent protein catabolic process |
| X | 1.8E+07 | G | A | CG6361 | 8.68E-05 | 0.047 | 0.049 | 0.013 | 0.044 | 1.58E-01 | serine-type endopeptidase activity;proteolysis;regulation of melanization defense response |
| X | 1.8E+07 | G | T | CG15042 | 2.20E-05 | 0.424 | 0.423 | 0.447 | 0.351 | 7.57E-01 | |
| X | 1.9E+07 | A | G | CCKLR-17D1 | 2.82E-05 | 0.078 | 0.061 | 0.050 | 0.016 | 1.49E-01 | G protein-coupled receptor activity;cholecystokinin receptor activity;integral component of plasma membrane;response to stress;G protein-coupled receptor signaling pathway;positive regulation of cytosolic calcium ion concentration;neuropeptide signaling pathway;neuromuscular junction development;neuropeptide receptor activity;adult locomotory behavior;larval locomotory behavior;gastrin receptor activity;integral component of membrane;neuronal cell body;terminal bouton |

| | | | | | | | | | | | |
|---|---|---|---|---|---|---|---|---|---|---|---|
| X | 2.1E+07 | T | C | CG34120 | 1.88E-05 | 0.059 | 0.086 | 0.094 | 0.056 | 5.49E-02 | transporter activity;ATP binding;transport;ATPase activity |
| X | 2.1E+07 | T | C | (CG32822) | 2.36E-05 | 0.044 | 0.031 | 0.015 | 0.050 | 3.46E-01 | NA |
| X | 2.2E+07 | C | T | (DIP1) | 7.27E-05 | 0.314 | 0.290 | 0.231 | 0.341 | NA | NA |
