## Supplemental Table 3 for "Microbiome composition shapes rapid genomic adaptation of *Drosophila melanogaster*"

| chrm | pos | ref | alt | gene | p.At_Lb | p.cli | concord | afMean .Founder | afMean .No-Ad | afMean .At | afMean .Lb | GO terms |
| --- | --- | --- | --- | --- | --- | --- | --- | --- | --- | --- | --- | --- |
| 2L | 9209955 | G | C | tai | 6.31E-03 | 9.50E-09 | FALSE | 0.033 | 0.034 | 0.021 | 0.052 | nucleus;regulation of transcription; DNA-templated;border follicle cell migration;nuclear receptor binding;nuclear receptor transcription coactivator activity;cellular response to hormone stimulus;nuclear hormone receptor binding;steroid hormone receptor binding;positive regulation of growth;positive regulation of transcription by RNA polymerase II;protein dimerization activity;axon extension;germ-line stem-cell niche homeostasis;positive regulation of border follicle cell migration |
| 2L | 14353549 | G | T | l(2)34Fd | 7.46E-03 | 3.53E-09 | TRUE | 0.159 | 0.209 | 0.247 | 0.185 | maturation of SSU-rRNA from tricistronic rRNA transcript;nucleolus;small-subunit processome |
| 2R | 738754 | G | A | (CG14464) | 6.47E-03 | 8.21E-12 | FALSE | 0.844 | 0.723 | 0.702 | 0.650 | NA |
| 3R | 10964114 | G | A | CG3984 | 2.28E-03 | 1.45E-09 | TRUE | 0.231 | 0.187 | 0.267 | 0.210 | |
| 3R | 12086524 | G | C | (tara) | 7.86E-03 | 4.97E-10 | FALSE | 0.342 | 0.363 | 0.326 | 0.402 | NA |
| 3R | 12254580 | G | T | (tRNA:CR31497) | 5.88E-03 | 1.21E-10 | TRUE | 0.073 | 0.115 | 0.121 | 0.047 | NA |

| | | | | | | | | | | | | |
|---|---|---|---|---|---|---|---|---|---|---|---|---|
| 3R | 12275145 | A | T | Pak3 | 4.01E-03 | 7.48E-10 | TRUE | 0.044 | 0.053 | 0.065 | 0.021 | ruffle;protein serine/threonine kinase activity;ATP binding;cytoplasm;protein phosphorylation;actin filament organization;myoblast fusion;cell migration;signal transduction by protein phosphorylation;stress-activated protein kinase signaling cascade;activation of protein kinase activity;protein homodimerization activity;regulation of MAPK cascade;protein autophosphorylation;Rac GTPase binding;regulation of axonogenesis;positive regulation of synapse assembly |

| | | | | | | | | | | | | |
|---|---|---|---|---|---|---|---|---|---|---|---|---|
| 3R | 12787654 | T | G | Abd-B | 1.04E-03 | 7.47E-09 | TRUE | 0.168 | 0.152 | 0.166 | 0.081 | transcription regulatory region sequence-specific DNA binding;RNA polymerase II distal enhancer sequence-specific DNA binding;pole cell migration;segment specification;specification of segmental identity; abdomen;open tracheal system development;salivary gland development;imaginal disc-derived genitalia development;midgut development;gonadal mesoderm development;heart development;sex differentiation;negative regulation of female receptivity;germ cell migration;male gonad development;negative regulation of cardioblast cell fate specification;male genitalia development;female genitalia development;genital disc development;genital disc anterior/posterior pattern formation;determination of genital disc primordium;external genitalia morphogenesis;genital disc sexually dimorphic development;spiracle morphogenesis; open tracheal system;negative regulation of salivary gland boundary specification;negative |
| 3R | 12811795 | T | C | CG18622 | 1.52E-03 | 6.40E-09 | TRUE | 0.027 | 0.046 | 0.064 | 0.033 | |
| 3R | 12835934 | T | C | CG8907 | 6.88E-03 | 3.48E-09 | TRUE | 0.444 | 0.403 | 0.439 | 0.319 | **actin binding**;plasma membrane;Rho protein signal transduction;Rac guanyl-nucleotide exchange factor activity;regulation of Rho protein signal transduction;olfactory behavior;synapse |

| | | | | | | | | | | | | |
|---|---|---|---|---|---|---|---|---|---|---|---|---|
| 3R | 12900510 | G | A | Keap1 | 9.85E-03 | 3.82E-13 | TRUE | 0.134 | 0.187 | 0.252 | 0.155 | **actin binding**;nucleus;polytene chromosome puff;response to oxidative stress;protein homodimerization activity;protein autoubiquitination |
| 3R | 12900525 | G | T | Keap1 | 2.81E-03 | 5.17E-13 | TRUE | 0.130 | 0.192 | 0.249 | 0.166 | **actin binding**;nucleus;polytene chromosome puff;response to oxidative stress;protein homodimerization activity;protein autoubiquitination |
| 3R | 12922911 | T | A | cher | 5.58E-03 | 4.38E-09 | TRUE | 0.692 | 0.696 | 0.615 | 0.685 | mitotic cell cycle;**actin binding**;actin filament;actin assembly;actin cytoskeleton;female germline ring canal formation;cytoplasmic transport;nurse cell to oocyte;long-term memory;motor neuron axon guidance;protein localization;determination of adult lifespan;Z disc;germarium-derived female germ-line cyst encapsulation;negative regulation of lamellocyte differentiation;apical cortex;sarcomere organization;behavioral response to ethanol;perinuclear region of cytoplasm;positive regulation of cytoskeleton organization;contractile ring |
| 3R | 12950403 | T | C | (CG31269) | 2.55E-03 | 1.50E-09 | TRUE | 0.030 | 0.055 | 0.072 | 0.024 | NA |
| 3R | 13636748 | A | G | CG18012 | 2.09E-03 | 1.70E-10 | TRUE | 0.074 | 0.065 | 0.094 | 0.046 | |
| 3R | 14009775 | C | T | CG7208 | 2.11E-04 | 5.70E-14 | TRUE | 0.064 | 0.068 | 0.073 | 0.026 | |
| 3R | 14193535 | C | T | Nup43 | 4.48E-03 | 4.42E-09 | TRUE | 0.074 | 0.086 | 0.111 | 0.035 | mitotic cell cycle;nuclear pore outer ring |

| | | | | | | | | | | | | |
|---|---|---|---|---|---|---|---|---|---|---|---|---|
| 3R | 14490730 | C | A | Gos28 | 5.01E-03 | 2.66E-10 | TRUE | 0.066 | 0.069 | 0.124 | 0.085 | Golgi trans cisterna;Golgi membrane;SNAP receptor activity;Golgi medial cisterna;cis-Golgi network;endoplasmic reticulum to Golgi vesicle-mediated transport;intra-Golgi vesicle-mediated transport;vesicle fusion;membrane;integral component of membrane;vesicle-mediated transport;SNARE complex;Golgi vesicle transport;regulation of vesicle targeting |
| 3R | 14647260 | T | G | CG18208 | 4.06E-03 | 3.99E-10 | TRUE | 0.091 | 0.135 | 0.162 | 0.119 | |
| 3R | 15874531 | T | A | CG17193 | 1.24E-03 | 4.96E-09 | TRUE | 0.220 | 0.258 | 0.339 | 0.239 | |
| 3R | 16514202 | T | A | (Oamb) | 3.77E-03 | 2.35E-09 | TRUE | 0.138 | 0.150 | 0.213 | 0.086 | NA |
| 3R | 17066761 | A | T | (e) | 3.25E-03 | 3.04E-11 | TRUE | 0.197 | 0.222 | 0.237 | 0.131 | NA |
| 3R | 17941912 | T | A | Ir94b | 6.93E-03 | 3.80E-09 | TRUE | 0.045 | 0.067 | 0.065 | 0.028 | detection of chemical stimulus;ligand-gated ion channel activity;integral component of membrane |

| | | | | | | | | | | | | |
|---|---|---|---|---|---|---|---|---|---|---|---|---|
| 3R | 19013520 | T | A | cnc | 8.34E-03 | 2.39E-09 | TRUE | 0.050 | 0.057 | 0.062 | 0.014 | DNA binding;DNA-binding transcription factor activity;nucleus;polytene chromosome puff;response to oxidative stress;oocyte dorsal/ventral axis specification;regulation of pole plasm oskar mRNA localization;blastoderm segmentation;oocyte microtubule cytoskeleton polarization;determination of adult lifespan;regulation of bicoid mRNA localization;response to endoplasmic reticulum stress;intestinal stem cell homeostasis;bicoid mRNA localization;pole plasm oskar mRNA localization;positive regulation of transcription by RNA polymerase II;protein heterodimerization activity;dendrite morphogenesis;oocyte nucleus localization involved in oocyte dorsal/ventral axis specification;head development;pharynx development |

| | | | | | | | | | | | | |
|---|---|---|---|---|---|---|---|---|---|---|---|---|
| 3R | 19696100 | T | A | Pli | 2.67E-03 | 7.07E-09 | TRUE | 0.059 | 0.078 | 0.111 | 0.046 | protein polyubiquitination;positive regulation of antimicrobial peptide production;ubiquitin-protein transferase activity;protein binding;cytosol;Toll signaling pathway;cytoplasmic side of plasma membrane;kinase regulator activity;innate immune response;negative regulation of Toll signaling pathway;positive regulation of Toll signaling pathway;ubiquitin protein ligase activity;protein K48-linked ubiquitination;negative regulation of defense response to bacterium;negative regulation of antifungal innate immune response |
| 3R | 19965402 | A | C | Npc2f | 1.01E-03 | 1.15E-12 | TRUE | 0.028 | 0.030 | 0.055 | 0.025 | sterol transport;intracellular cholesterol transport;sterol binding |
| 3R | 19979636 | C | T | CG6178 | 2.40E-03 | 4.06E-12 | TRUE | 0.039 | 0.062 | 0.078 | 0.026 | long-chain fatty acid metabolic process;fatty-acyl-CoA synthase activity;long-chain fatty acid-CoA ligase activity;peroxisome;fatty acid biosynthetic process;acyl-CoA metabolic process;fatty acid ligase activity;CoA-ligase activity;luciferin monooxygenase activity;fatty-acyl-CoA biosynthetic process |
| 3R | 19979912 | G | A | CG6178 | 2.16E-03 | 9.63E-11 | TRUE | 0.044 | 0.065 | 0.073 | 0.029 | long-chain fatty acid metabolic process;fatty-acyl-CoA synthase activity;long-chain fatty acid-CoA ligase activity;peroxisome;fatty acid biosynthetic process;acyl-CoA metabolic process;fatty acid ligase activity;CoA-ligase activity;luciferin monooxygenase activity;fatty-acyl-CoA biosynthetic process |
| 3R | 20077768 | A | G | CG5706 | 8.69E-03 | 1.59E-12 | TRUE | 0.076 | 0.096 | 0.097 | 0.054 | |

| | | | | | | | | | | | | |
|---|---|---|---|---|---|---|---|---|---|---|---|---|
| 3R | 20165522 | G | A | CG6432 | 6.77E-03 | 6.70E-11 | FALSE | 0.477 | 0.412 | 0.379 | 0.498 | catalytic activity;fatty acid biosynthetic process;short-chain fatty acid-CoA ligase activity |
| 3R | 20165558 | C | G | CG6432 | 2.18E-03 | 4.17E-11 | FALSE | 0.457 | 0.465 | 0.454 | 0.533 | catalytic activity;fatty acid biosynthetic process;short-chain fatty acid-CoA ligase activity |
| 3R | 20341694 | C | T | CG13617 | 6.59E-03 | 1.47E-09 | TRUE | 0.044 | 0.063 | 0.071 | 0.020 | nucleic acid binding;cytoplasm;ciliary basal body;cilium assembly |
| 3R | 20570027 | T | G | tok | 4.36E-03 | 1.74E-11 | TRUE | 0.246 | 0.347 | 0.350 | 0.211 | metalloendopeptidase activity;calcium ion binding;axon guidance;defasciculation of motor neuron axon;motor neuron axon guidance;zinc ion binding;imaginal disc-derived wing vein morphogenesis;negative regulation of gene expression;protein processing |
| 3R | 20653990 | G | T | (niki) | 5.51E-03 | 9.57E-11 | FALSE | 0.053 | 0.058 | 0.034 | 0.084 | NA |
| 3R | 22886860 | G | T | Nep5 | 3.52E-03 | 2.54E-09 | FALSE | 0.413 | 0.437 | 0.403 | 0.448 | metalloendopeptidase activity;proteolysis;metallopeptidase activity;integral component of membrane |
